# Locomotor activity enhances environmental representation by boosting visuo-frontal communication

**DOI:** 10.64898/2026.11.02.763783

**Authors:** Russell Milton, Mitchell Slapik, Sergey Egranov, Arun Parajuli, Melissa Franch, Alexandra McConnell, Valentin Dragoi

**Affiliations:** Department of Neurobiology and Anatomy McGovern Medical School University of Texas - Houston Houston, TX, USA; Center for Neural Systems Restoration Houston Methodist Research Institute Houston, TX, USA; Neuroengineering Initiative Rice University Houston, TX, USA; Brain and Mind Research Institute Department of Physiology and Biophysics Weill Cornell Medical College Cornell University New York, NY, USA

## Abstract

During environment exploration we rapidly acquire visual information to identify targets of interest. Here, we discovered that locomotion limits how long natural scenes are viewed during free exploration. Whether the brain has developed compensatory mechanisms to ensure that sensory information is accurately transmitted to downstream areas during and after locomotion is unknown. We examined this issue by recording the spiking activity of populations of neurons in visual and prefrontal cortex in conjunction with wireless recordings of oculomotor events while macaques moved freely to explore their environment or performed treadmill bipedal locomotion. The communication between visuo-frontal cortical areas was enhanced during and after locomotion to increase visual-evoked responses and population coding accuracy in prefrontal, but not visual cortical neurons. This in turn improved behavioral performance in a freely moving visual detection task after locomotion. Electrical stimulation further revealed elevated inter-areal synaptic efficacy during locomotor activity. The mechanism whereby locomotion enhances stimulus coding accuracy in prefrontal cortex and behavioral performance by increasing visuo-frontal communication is functionally beneficial during environmental exploration when incoming stimuli are rapidly changing and the window for perceptual decisions is temporally restricted.

---

While actively exploring our environment, we navigate between different spatial locations to visually acquire information and transmit it to higher brain areas where perceptual decisions are made. This process must occur very rapidly as each image is only viewed for a fraction of a second^1^. Here, we discovered that locomotor activity restricts the duration for which natural scenes are viewed during free exploration. That is, visual fixations during unrestrained environment exploration are shorter when preceded by active wakefulness compared with quiescence. This implies that transmitting visual information across brain areas may be particularly challenging during and after locomotion when decision-making areas must rely on temporally limited information. However, even though this process occurs thousands of times over the time course of a day during environment exploration, it remains unknown whether and how locomotion impacts neuronal interareal communication.

Did the brain develop compensatory mechanisms to ensure that stimulus information is accurately transmitted to downstream areas during active wakefulness when sensory cortical neurons process incoming stimuli for shorter periods of time relative to periods of quietness? We performed simultaneous multi-electrode recordings in visual area V4 and dorsolateral prefrontal cortex (dlPFC) to examine the environmental encoding and inter-areal communication while macaques freely explored their environment or performed bipedal treadmill locomotion. We hypothesized that the accuracy of transmitting visual information depends on the strength of synaptic connections between sensory and downstream cortical areas. Locomotion strengthened the coupling between visual and prefrontal decision-making areas, enhancing stimulus-evoked responses and coding accuracy in downstream cortical neurons. Furthermore, behavioral responses in a color change task in freely moving conditions were more accurate when visuo-frontal communication was increased after locomotion.

## RESULTS

### Locomotion selectively enhances prefrontal cortical responses during free exploration

We recorded the activity of populations of V4 and dlPFC neurons using chronically-implanted multi-electrode arrays in conjunction with monitoring oculomotor events while animals freely explored an arena (1,564 neurons in V4 and 2,206 neurons in dlPFC, n = 27 sessions in 2 animals, **Fig. 1a** and **Fig. 1b**) where 3D objects of various shapes, colors, and contours were positioned (see Methods). Locomotion increased pupil diameter (**Extended Data Fig. 1a**), which is consistent with elevated arousal (**Fig. 1d**). We analyzed the overhead video recorded to track animal’s movement in the arena and define epochs of “active” (animal moving inside the arena) and “quiet” wakefulness (animal sitting quietly, but visually engaged). This was done by creating a composite image of the arena background and then subtracting it from each frame reveal the animal’s outline: a centroid of this shape pinpointed the animal’s location, while changes in this location over time determined the animal’s motion. For each session, the median motion during the awake state was used as a threshold to classify each 5-s epoch as active or quiet wakefulness. We investigated how these states influence the coding of natural stimuli during free viewing across multiple cortical areas.

**Figure 1:**
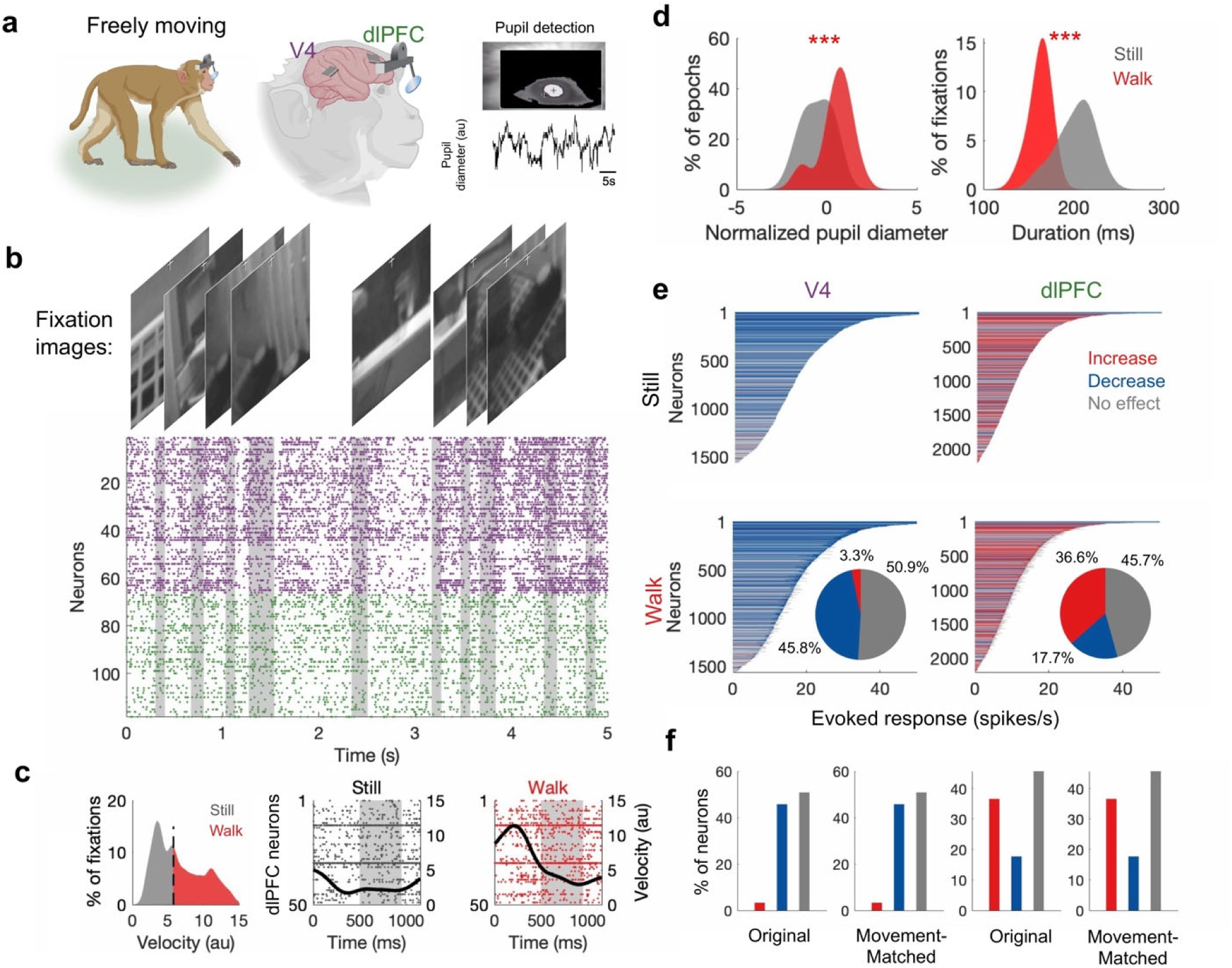
Naturalistic locomotion increases stimulus-evoked responses in dlPFC. **a,** Neural recordings from areas V4 and dlPFC in freely moving macaques with simultaneous tracking of visual input using wireless eye trackers. **b,** Visual input during fixations (vertical gray bars) aligned with neural activity across neurons in V4 (purple) and dlPFC (green). **c,** Left panel: Distribution of 5-s animal movement velocity preceding visual fixation for the still and walk conditions. Middle and right panels: Movement velocity as a function of time (thick black line) overlaid on example population rasters during still and walk conditions; vertical gray window represents the fixation period. **d,** Distribution of pupil diameters during still and walk epochs (left); distribution of fixation durations during still and walk epochs. **e,** Neural activity during the still and walk conditions in V4 (12.1% decrease, Wilcoxon rank-sum test; p = 5.84 × 10^-168^, n = 1,564 neurons) and dlPFC (4.25% increase; Wilcoxon rank-sum test; p = 2.38 × 10^-40^, n = 2,206 neurons). Inset shows proportion of neurons that increased, decreased, or maintained their firing rates during walk condition. **f,** Original and movement-matched stimulus-responses between still and walk conditions. Red represents neurons that significantly increased their firing rates during walk, blue represents neurons that decrease their firing rates, and gray represents neurons with no significant change. * p < 0.05. ** p < 0.01, *** p < 0.001.

We extracted thousands of fixation episodes (average of 3,612 fixations per session) and recorded the stimuli inside each cell’s receptive field. Then we assigned each fixation to a still and walk period based on the mean velocity of movement in the preceding 5-s interval (**Fig. 1c**). Pupil diameter was significantly higher during walk compared to the still condition (Wilcoxon signed-rank test; p = 2.23 × 10^-5^), but there was no difference in eye velocity (Wilcoxon signed-rank test, p = 0.79; X-position variance (two-sample t-test, p = 0.66) or Y-position variance (two-sample t-test, p = 0.94) (**Extended Data Fig. 1**). The scenes viewed during the still and walk periods did not significantly differ in mean brightness (Wilcoxon signed-rank test, p = 0.57, n = 27 sessions), or mean contrast (Wilcoxon signed-rank test, p = 0.40, n = 27 sessions) across 97,406 total frames. Importantly, to rule out the effects of movement on neural activity, we only analyzed stimulus responses during visual fixations when the animal was stationary, where each fixation was preceded by 5 seconds of still or walk. Across sessions, fixation durations were significantly shorter in the walk compared with still condition (two-sample t-test, 3.28 × 10^-12^), hence limiting the time window for gathering sensory information after locomotion (**Fig. 1d**). Simultaneously, locomotion had a significant effect on stimulus-evoked responses in both cortical areas. Specifically, during fixations preceded by active exploration, responses were decreased in V4 (Wilcoxon rank-sum test; p = 5.84 × 10^-168^, n = 1,564 neurons). Proportionally, 45.78% of V4 neurons showed a significant decrease while 3.32% of V4 neurons showed a significant increase in responses (**Fig. 1e**; see also **Extended Data Fig. 2** for example trials). These results are somewhat surprising as previous mouse studies had reported an almost doubling of neuronal responses in V1 during locomotion at similar speeds^2,3^. Meanwhile, we found the opposite results in dlPFC. During fixations preceded by active exploration, responses were significantly increased (Wilcoxon rank-sum test; p = 2.38 × 10^-40^, n = 2,206 neurons) (**Fig. 1e)**. Proportionally, locomotion modulated a substantial portion of dlPFC neurons: 36.58% showed a significant increase, 17.72% showed a significant decrease.

Even though we only examined neural responses to visual fixations when the animal was stationary, it has been previously shown that small body movements are correlated with dlPFC activity^4^. To eliminate this possible confound, we measured the degree of body movement during each fixation by quantifying the pixel changes in the view of the overhead camera. Although, on average, body movements were greater during walk relative to the still condition (**Extended Data Fig. 3;** one-way t-test, p = 3.10 × 10^-11^), we selected a subset of fixations wherein body movements were not significantly different between conditions (one-way t-test, p = 0.86; this only excluded approximately 10% of total fixations). However, our main results remained unchanged (**Extended Data Fig. 4a**) – locomotion significantly increased stimulus-evoked responses in dlPFC (Wilcoxon rank-sum test; p = 4.12 × 10^-27^; n = 1,979 neurons: 34.61% showed a significant increase in responses while 17.18% of cells showed a significant decrease), but V4 responses were decreased (Wilcoxon rank-sum test; p = 1.43 × 10^-151^, n = 1,395 neurons, **Fig. 1f**; 47.03% of V4 neurons showed a significant response decrease and 3.66% of neurons showed a significant increase during locomotion).

### Bipedal treadmill locomotion

Further, we recorded the activity of populations of V4 and dlPFC neurons as head-fixed macaques viewed stimuli on a computer screen while standing still or walking on a treadmill (**Fig. 2a**, see Methods; 628 neurons in V4 and 389 neurons in dlPFC). To calibrate the speed of the treadmill, we examined 60-min video recordings of monkeys freely walking in an enclosure^5^ (see Methods). On average, animals moved at 5.43 cm/s, although they routinely reached higher speeds. Therefore, we chose a treadmill belt speed of 8 cm/s, as this was well within the range of normally observed speeds, yet slow enough to maintain stable recordings. Bipedal locomotion increased general arousal^5–8^, i.e., pupil diameter was significantly elevated during periods of walking (**Fig. 2c**; Wilcoxon signed-rank test, p = 5.58 × 10^-7^, n=22 sessions; doubling treadmill speed did not further increase pupil size, **Fig. 2c**; Wilcoxon rank sum p = 0.29). Additional tests confirmed that there was no significant difference in eye velocity between the still and walk conditions (**Extended Data Fig. 5**, Wilcoxon signed-rank test, p = 0.83, X eye position variance, two-sample t-test, p = 0.24; Y eye position variance, two-sample t-test, p = 0.37).

**Figure 2:**
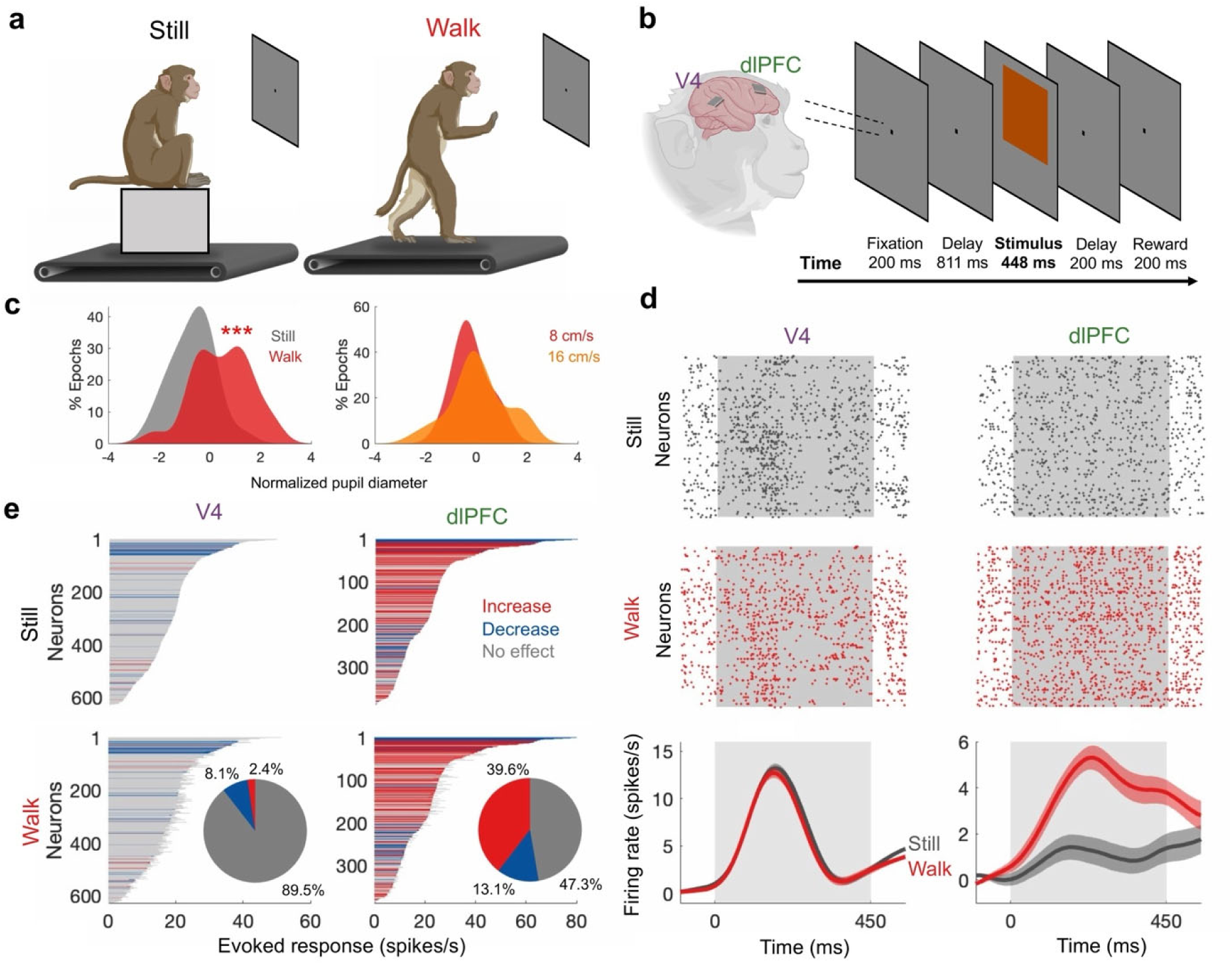
Bipedal treadmill locomotion increases stimulus-evoked responses in dlPFC but not V4. **a,** Animals performed a visual fixation task under both still (left) and walk (right) conditions. **b,** Visual task consisted of: (1) eye fixation, (2) stimulus presentation (color patch) and (3) reward at fixed time intervals since stimulus offset. **c,** Left panel: Continuous tracking of pupil size and location with ISCAN eye-tracker. Center panel: Distribution of pupil diameter during the still and walk conditions (Wilcoxon signed-rank test, p = 6.98 × 10^-5^, n = 22 sessions). Right panel: Distribution of pupil diameter between the two different walking speeds, fast and slow (Wilcoxon signed-rank test, p = 0.29, n = 31 sessions). **d,** Top and middle rows: Example population rasters in V4 (left) and dlPFC (right) for still and walk conditions. Gray shaded region denotes the time of stimulus presentation. Bottom row: Population firing rates averaged across all stimulus-responsive neurons in V4 and dlPFC. Baseline firing rates have been subtracted. Traces show mean +/- SEM. **e,** Stimulus responses in dlPFC (increase of 11.72%; Wilcoxon rank-sum test; p = 5.76 × 10^-15^, n = 389 neurons) and V4 (Wilcoxon rank-sum test; p = 0.83, n = 628 neurons). Inset shows proportion of neurons that increase, decrease, or maintain their firing rates during locomotion. * p < 0.05. ** p < 0.01, *** p < 0.001.

Monkeys performed a visual fixation task in the still and walk conditions while 8 isoluminant color patches with equal separation in the CIELUV perceptually-uniform color space were briefly flashed on the screen for 448 ms across neurons’ receptive fields^9^ (**Fig. 2b**, see Methods; recordings in the 2 conditions were performed in a blocked manner). Across the population of cells, treadmill locomotion modulated V4 responses to visual stimuli only weakly. Overall, locomotion yielded no significant effect on stimulus-evoked V4 responses (**Fig. 2d**, Wilcoxon rank-sum test; p = 0.83). Proportionally, the locomotion-associated modulation of individual neuron responses was statistically significant in only a minority of units (**Fig. 2e**), i.e., 2.39% of V4 units showed a significant increase, and 8.12% showed a significant decrease in responses (false-discovery rate, FDR-adjusted p-values < 0.05; n = 628 units).

Overall, while V4 neurons showed no effect of locomotion, dlPFC responses were significantly modulated (**Fig. 2e**; 11.72% increase; Wilcoxon rank-sum test; p = 5.76 × 10^-15^, n = 389 cells). Comparing neurons between the still and walk conditions, we found that locomotion significantly increased stimulus responses in 39.59% of units, while 13.11% of units showed a significant decrease (**Fig. 2e**; see also **Extended Data Fig. 6** for example trials and **Extended Data Fig. 7** for an analysis of ongoing activity before stimulus presentation). Locomotion did not influence stimulus selectivity (**Extended Data Fig. 8**) or response latency (**Extended Data Fig. 9**). To control for possible changes in ongoing firing rates across conditions, we repeated the above analysis by selecting the neurons in both areas that did not change their baseline firing between the still and walk conditions (**Extended Data Fig. 10a)**. However, even for this subpopulation, locomotion significantly increased stimulus responses in dlPFC (4.47% increase, Wilcoxon signed-rank test: p = 5.00 × 10^-6^, n = 190 neurons), while responses in V4 were only slightly modulated by walking (0.86% increase, Wilcoxon signed-rank test: p = 0.033, n = 559 neurons).

### Locomotion boosts inter-areal communication

We hypothesized that the elevated responses during locomotion in prefrontal cortex could be due to enhanced communication between visuo-frontal areas. Using linear regression, we examined the impact of changes in brain state induced by locomotion on the interaction between neurons with overlapping receptive fields in the same or different areas. Cortical areas generate diverse patterns of activity, but only some of these patterns are passed on through inter-areal communication. This mechanism of sharing some activity patterns but not others based on “communication subspaces” has been hypothesized to selectively send information to specific areas for further processing^10^. To calculate subspace communication, we computed the residual visual responses of each neuron by subtracting its mean response to obtain trial-by-trial fluctuations around the mean (**Fig. 3a**), and then used a linear regression on the fluctuations in one population to predict the fluctuations in another (**Fig. 3b**). The accuracy of this prediction provides a measure of intra or inter-areal communication. During the still epochs of the freely moving experiment, we predicted 30.85% and 17.83% of variance within areas V4 and dlPFC, and 6% for feedforward and 6.18% for feedback communication respectively (**Fig. 3c**, where dimensions = 10).

**Figure 3:**
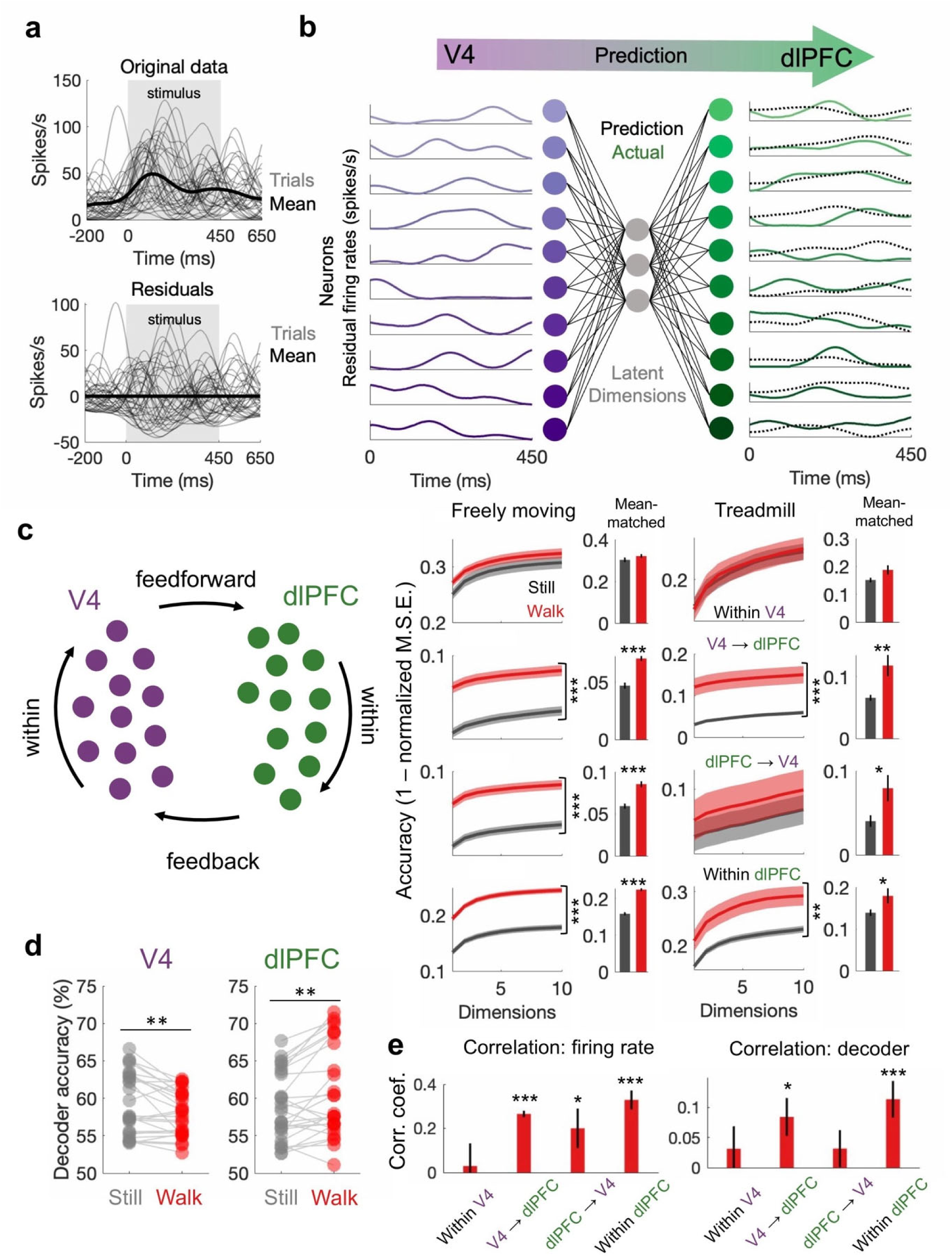
Locomotion increases visuo-frontal communication in both treadmill and freely-moving experiments. **a,** Stimulus responses and residual fluctuations for an example V4 neuron. **b,** Example reduced-rank linear regression predicting response fluctuations in dlPFC (green) based on fluctuations in V4 (purple). **c,** Subspace communication within and between cortical areas for the freely-moving (left) and treadmill (right) experiments. Traces show mean +/- SEM. Bar graphs with error bars (SEM) show subspace communication for mean-matched neurons that did not change their ongoing firing rates between the Still and Walk conditions. **d,** Decoding of stimulus identity in the freely-moving experiment for V4 (paired t-test, p = 0.0038, n = 27 sessions) and dlPFC (paired t-test, p = 0.0018, n = 27 sessions). Lines connect dots from the same session. **e,** Left panel: Correlation between the change in subspace weight and change in firing rate between the Still and Walk conditions. Right panel: Correlation between change in subspace weights and the change in decoder weights between the Still and Walk conditions. * p < 0.05. ** p < 0.01, *** p < 0.001.

Locomotion significantly modulated communication within and between cortical areas: in feedforward direction, a 48.38% increase, two-sample t-test, p = 7.84 × 10^-8^, n = 270 folds, the feedback direction (46.18% increase, two-sample t-test, p = 2.54 × 10^-9^). and within dlPFC, a 37.20% increase, two-sample t-test, p = 6.80 × 10^-34^, n = 270 folds, but not V4 (two-sample t-test, p = 0.19, n = 270 folds). Similarly, during the still epochs of the treadmill experiment, we predicted 23.3% and 23.0% of variance within areas V4 and dlPFC, respectively (**Fig. 3c**, where dimensions = 10) while across areas we predicted a smaller but significant proportion of the variance: 5.9% for feedforward (V4 to dlPFC) and 8.4% for feedback communication (dlPFC to V4). During bipedal treadmill locomotion, neuronal communication was improved in the feedforward direction (153.25% increase, two-sample t-test, p = 0.0015, n = 130 folds) and within dlPFC (22.61% increase, two-sample t-test, p = 2.41 × 10^-5^, n = 130 folds). In contrast, locomotion did not modulate communication between V4 neurons (two-sample t-test, p = 0.83, n = 130 folds) or feedback communication (two-sample t-test, p = 0.43, n = 130 folds), possibly due to the more passive nature of viewing in head-fixed experiments. Further examining neuronal communication using a second measure, spike-timing coordination, revealed that locomotion significantly increased spike-timing coordination between visuo-frontal cortical areas and within dlPFC (**Extended Data Fig. 11**, see Methods).

The improved inter-areal communication during locomotion could result from a decrease in the firing threshold of dlPFC neurons, but not V4 neurons. To control for this possibility, we examined the subspace communication between the neurons that did not change their ongoing firing rates between the still and walk conditions. Locomotion still improved inter-areal communication in feedforward (50.28% increase, two-sample t-test, 7.66 × 10^-12^, n = 270) and feedback directions (43.98% increase, two-sample t-test, p = 2.50 × 10^-8^, n = 270 folds), as well as within dlPFC (41.50% increase, two-sample t-test, p = 8.87 × 10^-29^, n = 270 folds), but not within V4 (two-sample t-test, p = 0.208, n = 270 folds). Performing the same control analysis on neurons that did not change their ongoing firing rate during the treadmill experiment, locomotion still improved inter-areal communication for feedforward (increase of 79.39%, two-sample t-test, p = 0.0036, n = 130 folds) and feedback (increase of 97.25%, two-sample t-test, p = 0.023, n = 130 folds), and communication within dlPFC (29.1% increase, two-sample t-test, p = 0.0397, n = 130 folds, **Fig. 3c** bar graphs, **Extended Data Fig. 6b**), but not V4 (two-sample t-test, p = 0.0513, n = 130 folds). Thus, improved inter-areal communication reflects elevated information transmission, and not simply a lower firing threshold of dlPFC neurons during locomotor activity.

Could body movements account for changes in inter-areal communication during locomotion? To address this issue, we used a subset of fixations eliciting responses that were ‘mean matched’ for body movements between still and walk conditions (**Extended Data Fig. 3**). However, even under these conditions our results remained unchanged – locomotion improved feedforward (**Extended Data Fig. 4b;** 48.51% increase, two-sample t-test, p = 4.94 × 10^-7^, n = 250) and feedback communication (46.65% increase, two-sample t-test, p = 1.40 × 10^-9^, n = 250), while also improving neuronal communication within dlPFC (31.68% increase, two-sample t-test, p = 4.63 × 10^-14^, n = 250 folds), but not within V4 (two-sample t-test, p = 0.19, n = 250 folds). Thus, body movements cannot explain the changes observed in subspace communication between the still and walk conditions.

### Locomotion improves coding accuracy in dlPFC but not V4

Does improved inter-areal communication during locomotion lead to improved encoding of the environment? Given the richness of visual information during environment exploration, stimulus identity was decoded from neural population activity in each cortical area during individual fixations in the freely-moving condition. An autoencoder condensed snapshots from the scene camera into a short image code, and then we created clusters of images that had similar image codes (see Methods); each cluster contained 150 fixations. Finally, an SVM linear decoder distinguished between two image clusters based on the pattern of neural activity during the corresponding fixation window, and this analysis was repeated for 200,000 cluster pairs. Overall, the walk condition was associated with enhanced stimulus coding in dlPFC (paired t-test, p = 0.0018, n = 27 sessions), but decreased coding accuracy in V4 (**Fig. 3d, Extended Data Fig. 12**, paired t-test, p = 0.0038, n = 27 sessions). Repeating this analysis for the treadmill experiment yielded consistent results: locomotion significantly increased decoder accuracy in dlPFC (p = 0.03, n = 130 folds), but not V4 (p = 0.010, n = 130 folds).

We further examined whether the effects of locomotion, i.e., changes in neurons’ firing rates, strength of inter-areal communication, and population coding accuracy occur in the same population of cells, or they occur independently of each other. We thus created a summary subspace weight for each neuron by averaging all the weights associated with its inputs. We also computed each neuron’s change in mean firing rate and mean decoder weight across the still and walk conditions. We found a significant correlation between the change in subspace weight and the change in firing rate between still and walk conditions (**Fig. 3e**) for feedforward (one-sample t-test, R = 0.27, p = 2.34 × 10^-16^) and feedback communication (one-sample t-test, R = 0.20, p = 0.034), and for communication within dlPFC (one-sample t-test, R = 0.33, 3.81 × 10^-8^), but not V4 (one-sample t-test, P = 0.78). The fact that increased subspace communication and increased firing rates are highly correlated indicates that inter-areal communication accounts for the higher firing rates in dlPFC in the walk condition. Similarly, we examined the change in subspace weights and decoder weights for each neuron between still and walk (**Fig. 3e**). We found a significant positive correlation between changes in subspace weights and decoder weights for feedforward (one-sample t-test, R = 0.084, p = 0.013) and dlPFC communication (one-sample t-test, R = 0.11, p = 9.30 × 10^-4^), but not V4 (one-sample t-test, p = 0.40) or feedback communication (one-sample t-test, p = 0.31). Thus, improved neuronal communication during locomotion increases both the firing rates of neurons and stimulus coding in the same dlPFC population.

### Locomotor activity improves behavioral performance

We further examined whether the elevated neuronal communication during locomotion enhances behavioral performance in a freely moving task (n = 8 sessions in 2 animals). Animals were placed in an arena with two reward stations spaced 120 cm apart and could freely navigate between them. Each station dispensed food reward based on a variable interval schedule: a red screen indicated that reward was available, while a blue screen indicated that it was not. Thus, animals had to use visual cues to determine when and where to press a button to receive a reward (**Fig. 4a**). Notably, because the stations were positioned 120 cm apart, animals often walked between the two sides of the arena, enabling us to divide the sessions into still and walk epochs based on when they switched sides. Consistent with our previous results (**Fig. 3)**, locomotion significantly improved interareal communication in the feedforward (Wilcoxon sign-rank test, p = 4.96 × 10^-4^, n = 100 folds) and feedback (Wilcoxon signed-rank test, p = 0.043, n = 100 folds) directions (**Fig. 4b**). Furthermore, locomotion improved behavioral performance as measured by push accuracy, i.e., the probability that a button push occurring in a particular state, still or walk, was rewarded (we excluded responses immediately before and after switching reward stations to eliminate potential confounds). After locomotion, animals pressed more accurately to obtain reward as the screen transitioned from blue to red (Wilcoxon signed-rank test, p = 0.0020, n = 10 sessions) (**Fig. 4c**). Across trials, we found a significant correlation between interareal communication and push accuracy (using bins of 10 consecutive button pushes, p = 0.049, n = 52 bins). Altogether, these results demonstrate that locomotion increases visuo-frontal communication to enhance behavioral performance.

**Figure 4:**
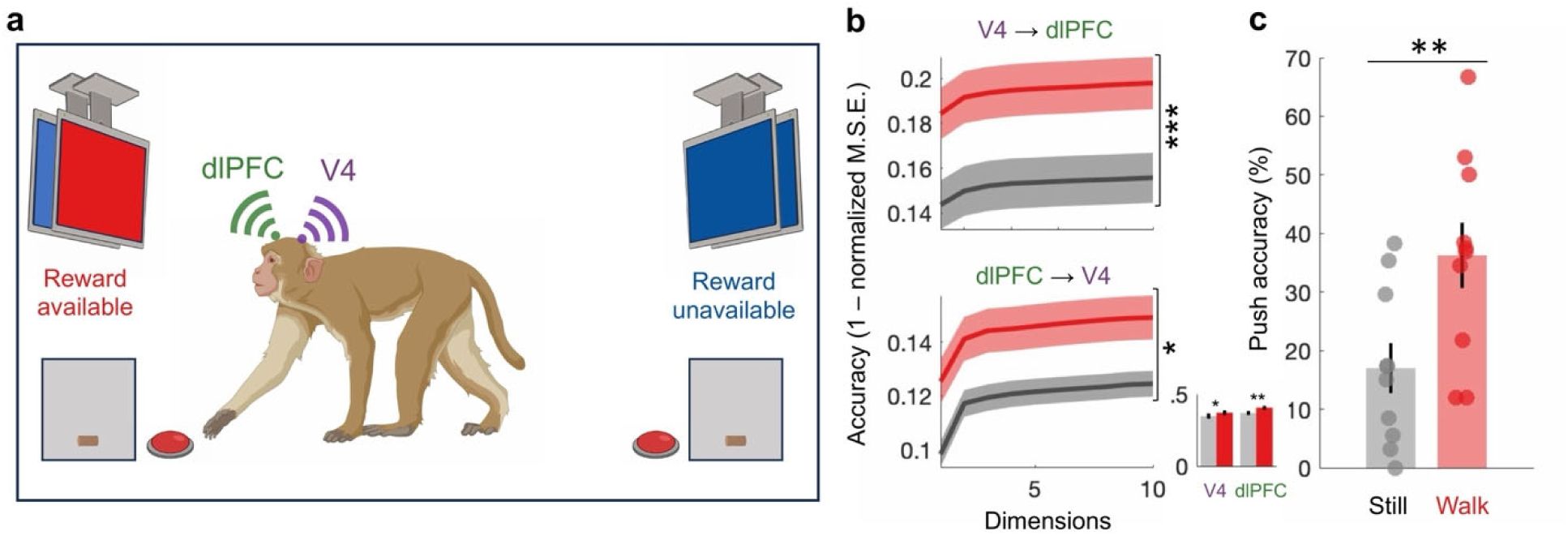
Locomotion improves behavioral performance in a freely moving color change task. **a,** Schematic of the task with two food stations, consisting of a monitor, button and food dispenser. Animals walked from one station to the other to receive reward. Shifting screen color from blue to red indicates that food is available after the next button push; blue screen indicates that food is unavailable. **b,** Subspace communication for still and walk epochs, measured as prediction accuracy between V4 and dlPFC (in both directions) for subspaces with different numbers of dimensions. **c,** Push accuracies during still and walk epochs. Accuracy measures the percentage of rewarded button pushes. * p < 0.05. ** p < 0.01, *** p < 0.001.

### Locomotion increases visuo-frontal synaptic efficacy

Finally, we used electrical stimulation to causally test whether locomotion improves inter-areal communication. This was done by developing an electrical stimulation (E-stim) assay involving stimulation of individual channels in one cortical area and simultaneously recording from distant postsynaptic cells in another area while head-fixed animals fixated on a blank screen (**Fig. 5a**). This allowed us to causally probe the strength of inter-areal synaptic efficacy when animals were either standing still or walking on the treadmill. On each electrode contact, we delivered repetitive 5 biphasic pulses (100 mA amplitude, 0.2 ms duration) at 50 Hz during a 500 ms fixation, followed by a 5-s delay (n = 7 sessions). The current amplitude was chosen to be at threshold level, i.e., the maximum current amplitude for which stimulation in one area did not increase responses in the postsynaptic area during the still condition. Yet during the walk condition, electrical pulses in V4 did elicit statistically significant responses in dlPFC based on a cross-correlation analysis (**Fig. 5b**, additional examples in **Extended Data Fig. 13**). Overall, 26.25% of the V4 electrodes significantly modulated firing rates in dlPFC across E-stim sessions. Next, we examined how locomotion modulated the propagation of these electrical stimulation-evoked signals. We found that electrical signals from V4 propagated more effectively to dlPFC during walk relative to the still condition, i.e., electrical stimulation in V4 increased dlPFC mean evoked responses during locomotion specifically during the 100-ms electrical stimulation epoch; two-sample t-test, p = 5.70 × 10^-10^, n = 2,456 trials). This indicates that locomotion enhances the feedforward propagation of information by increasing visuo-frontal synaptic efficacy (**Fig. 5c**).

**Figure 5:**
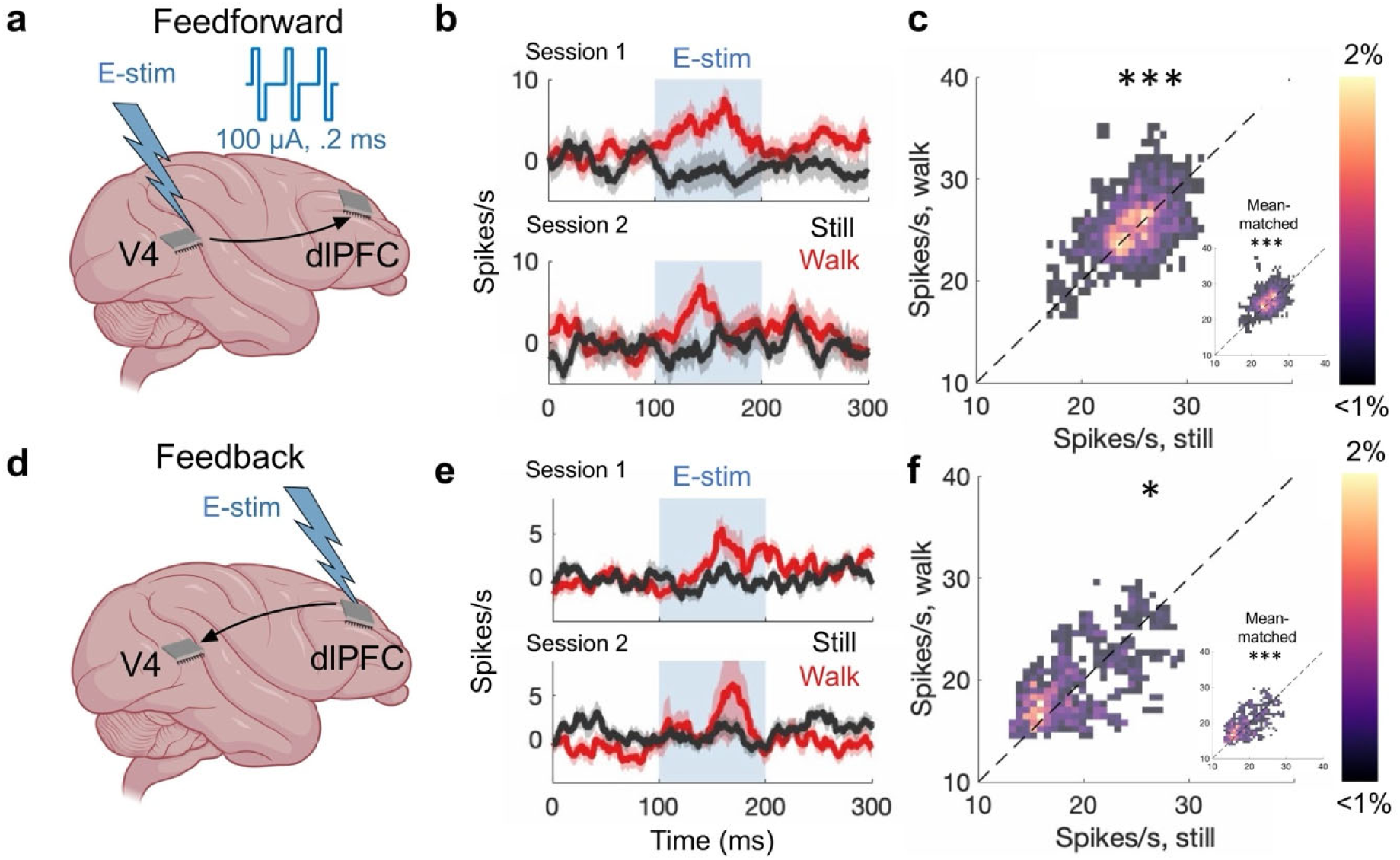
Electrical stimulation assay to measure changes in visuo-frontal synaptic efficacy during locomotion. **a,** Schematic of V4 electrical stimulation pulses propagating to dlPFC. **b,** Example dlPFC evoked responses (70 neurons) to electrical stimulation in V4. Traces show mean +/- SEM. **c,** dlPFC responses to V4 stimulation during the Still and Walk conditions (two-sample t-test, p = 5.70 × 10^-10^, n = 2,456 trials). Inset: dlPFC responses to V4 stimulation for mean-matched neurons that did not change their ongoing firing rate between the still and walk conditions (two-sample t-test, p = 1.30 × 10^-16^). **d,** Schematic of dlPFC electrical stimulation propagating to V4. **e,** Example V4 evoked responses (68 neurons) to electrical stimulation in dlPFC. Traces show mean +/- SEM. **f,** V4 evoked responses to dlPFC electrical stimulation during Still and Walk conditions (two-sample t-test, p = 0.011, n = 910 trials). Inset: V4 responses to dlPFC stimulation for ‘mean-matched’ neurons that did not change their ongoing firing rates between Still and Walk conditions (two-sample t-test, p = 9.99 × 10^-4^, n = 938 trials). In these probability distributions, yellow represents a high proportion of neuron pairs while black represents a low proportion. * = p < 0.05. ** = p < 0.01, *** = p < 0.001.

In principle, postsynaptic response enhancement could be attributed to a lower firing threshold in dlPFC rather than an enhanced communication between cortical areas. Thus, we further examined the effects of V4 stimulation on those dlPFC neurons that did not change their ongoing firing rate between the still and walk conditions. However, for this neuron subpopulation (n=207 cells), we still found that electrical stimulation in V4 evokes higher responses in dlPFC during walk condition relative to the still condition (two-sample t-test, p = 1.30 × 10^-16^, n = 2,155 trials). This indicates that the stronger evoked responses in dlPFC during locomotion are due to enhanced communication between cortical areas and not simply due to a lower firing threshold in dlPFC (**Fig. 5c**, inset).

The increased synaptic efficacy between visual and prefrontal cortex during locomotion extended beyond feedforward communication. When we electrically stimulated dlPFC neurons to evoke responses in V4 (**Fig. 5d**), we found that 35.38% of electrodes significantly modulated firing rates in V4 (**Fig. 5e**; additional examples in **Extended Data Fig. 13**), and these signals propagated more effectively to V4 during walk relative to the still condition (**Fig. 5f**; two-sample t-test, p = 0.011, n = 910 trials). Repeating this analysis for the V4 neurons that did not change their ongoing firing rates between the still and walk conditions still yielded a significant enhancement of feedback propagation of information (**Fig. 5f** inset; two-sample t-test, p = 9.99 × 10^-4^, n = 938 trials). Finally, we tested whether these results could be attributed to stronger evoked neuronal responses at the electrical stimulation site, either in V4 or dlPFC, during locomotion rather than increased visuo-frontal communication. Therefore, given that we cannot simultaneously electrically stimulate and record neuronal activity, we analyzed the mean spiking rate of each neuron at the stimulation site in the interval beginning 7 ms after each stimulation pulse (to avoid stimulation artifacts) and ending at the time of the next pulse, for each train of stimulation pulses. Overall, we found that locomotion did not influence the stimulation-evoked mean firing rates of neurons at the electrical stimulation site (Wilcoxon signed-rank test, p > 0.05 in V4 and dlPFC, comparing the still and walk conditions). Thus, increased signal propagation during locomotion must be due to enhanced interareal communication, rather than increased responses at the stimulation site.

## DISCUSSION

Although locomotion represents such an integral component of our daily activities, whether and how it impacts the environmental representation during perception and the communication between cortical areas and has been unknown. This issue is particularly important during environment exploration when visual fixations are brief, and the visual system must operate in a highly efficient manner. Does the brain possess compensatory mechanisms to ensure that sensory information is accurately transmitted to downstream areas during and after locomotion despite the fact that this information is available for shorter periods of time relative to quiet wakefulness? Using a novel approach combining bipedal treadmill locomotion, wireless recording of cortical population activity, and wireless eye tracking during free environment exploration, we found that locomotion enhances the communication between visuo-frontal cortical areas to increase visual-evoked responses and improve the representation of the environment in prefrontal cortex. This mechanism is particularly beneficial during active exploration when visual information about ethological stimuli changes rapidly, and perceptual decisions must be made on the fly.

Our results also reflect evolutionary differences in how locomotion modulates sensory representations in neocortex. High acuity perception in primates, including humans, is mediated by the fovea, and therefore primates extract visual information by making rapid fixational eye movements^1,11^. In contrast, rodents acquire visual information by making head movements, and eye movements have a compensatory role by stabilizing the visual scene^12^. As demonstrated here (**Fig. 1d**), locomotion reduces the duration of fixations in unrestrained non-human primates. Thus, the significant increase in visuo-frontal communication represents a compensatory mechanism by which prefrontal neurons increase their firing despite the decrease of responses in visual cortex. This differs from the previous studies in rodents indicating that locomotion shifts neural activity to a more desynchronized, heightened-arousal state, associated with increased visual-evoked responses and selectivity^13^. However, rodents are nocturnal prey animals with low visual acuity that spend most of their time in darkness. In contrast, primates generally spend their waking hours in well-lit conditions. Their high visual acuity enables them to extract salient information from distant objects, and primates have evolved specialized visual cortical areas to represent the environment in a hierarchical manner^5^. Furthermore, there are substantial differences between visuo-motor circuitry of rodents and primates^14^, which raises the possibility that rodent and primate brain might have evolved to employ different mechanisms by which locomotion influences how the environment in represented in the brain.

The increase in visuo-frontal synaptic efficacy during locomotion could reflect an arousal-mediated effect of neuromodulators on synaptic plasticity^15^. Locomotor effects on visual responses are likely to originate from basal forebrain projections targeting multiple areas throughout neocortex^16,17^. The differences across areas reported here may be due to region-specific differences in cholinergic circuits^13^. Future work will further our understanding of the functional implications of these distinct cholinergic circuits between sensory and executive cortical areas during locomotor activity and elucidate their impact on visuo-frontal synaptic communication.

## METHODS

### Ethics statement and animals

All experiments were performed in accordance with protocols approved by the Animal Welfare Committee (AWC) and the Institutional Animal Care and Use Committee (IACUC) for the University of Texas Health Science Center at Houston (UTHealth). Two adult male rhesus monkeys (Macaca mulatta; M: 10 kg, 11 years old; R: 12 kg, 10 years old) were used in the experiments.

### Chronic implantation of the Utah array

A titanium headpost was implanted medially with anchor screws on each monkey. After acclimatization, animals were implanted with a 64-channel dual Utah array in the left hemisphere dlPFC (anterior of the arcuate sulcus and dorsal of the principal sulcus), left V4 (ventrally anterior to lunate sulcus and posterior to superior temporal sulcus), and a pedestal on the caudal skull (Blackrock Neurotech). We used Brainsight, a neuronavigational system, and animal’s MRIs to determine the location for V4 and dlPFC craniotomies (Rogue Research). During surgery, visual identification of arcuate and principal sulci guided precise implantation of arrays into the dlPFC, and visual identification of the lunate and superior temporal sulci supported array placement in area V4. The dura was sutured over each array, and two reference wires were placed above the dura mater and under the bone flap. The bone flaps were secured over the arrays using titanium bridges and screws, and the electrical contacts on the pedestal were protected using a plastic cap except during the experiment. Following the implantation, the monkeys had a 3-week recovery period before recording from the arrays.

### Freely-moving electrophysiology

In the freely-moving experiment, we aimed to record the activity of neurons while minimizing interference with the animals’ behavior. Therefore, we used a lightweight, rechargeable battery-powered device (Cereplex-W, Blackrock Neurotech) that communicates wirelessly with a central amplifier and digital processor (Cerebus Neural Signal Processor, Blackrock Neurotech)^11,18^. In each session, neural activity was recorded in the head-fixed position for 10 min to ensure the signal quality before releasing the monkey in the experimental arena. The arena was surrounded by eight antennas. Spike-sorting was performed as described for head-fixed electrophysiology, above.

### Freely-moving eye tracking

In the freely-moving experiment, we used a custom wireless eye tracker (ISCAN) to measure pupil position and diameter as described previously. This portable eye tracker was mounted dorsally, right above the animal’s head, and consisted of an eye mirror, an eye camera, and a scene camera, which records animal’s field of view. Here, all data was recorded at 30 Hz. To train animals to wear the device without damaging it, we first 3D-printed model versions with Sketchup Pro and fitted them with eye mirrors. We also created custom adaptors to anchor the models to the animals’ headposts, without directly touching the animals to minimize discomfort and reduce tampering. Animals wore these model eye trackers for several mock recording sessions to familiarize them to wearing the device. Once they grew accustomed to wearing the model and stopped touching it altogether, it was swapped for the real device. Before each experiment, the eye tracker was secured to the animal and we performed a calibration procedure (‘point-of-regard’ calibration) while the animal was head-fixed, which mapped the eye position on the view of the head-mounted scene camera. Animals were trained to view five calibration points within the field of view of the scene camera (640 × 480 pixels or 35 × 28°): a center calibration point, and four outer points positioned at ±8 to ±10° with respect to the center; these distances were chosen based on the approximate range of stimuli distances during free viewing (70 cm). As the animal viewed each point, the eye-calibration software synchronized the eye-movement data with the image frames recorded by the scene camera. After calibration, the animal’s center of gaze was displayed on each scene camera frame in real time as crosshairs. If the animal looks outside the field of view of the scene camera, gaze location was not detected and the crosshairs did not appear on the frames. When this occurred, eye position was reported as zero. Therefore, we only used scene camera frames that included crosshairs, which made up the majority of frames. We used the horizontal and vertical coordinates of the pupil to compute eye speed. To extract fixations, we determined a conservative speed threshold that best separated the bimodal distribution of eye speed in each session. A fixation was defined as a minimum of 100 ms when the eye speed remained below this threshold.

### Movement quantification

Unlike traditional eye tracking systems mounted to an external structure, our wireless eye tracking system for free roaming monkey was directly mounted to the monkey’s head. For this reason, there are rare instances when the monkey’s gaze direction remains constant even while his head and field of view are moving. To prevent these moments from being false identified as fixations, we estimated the motion of the scene content between two consecutive frames (*‘pixel-flow’*) of the scene camera video using the "BlockMatcher" method available in the Matlab Vision toolbox. *Pixel-flow* is a measure that estimates how much the content of a frame has moved relative to its position in the reference frame by moving a block of pixels over a search region and computing their similarities. We quantified the *pixel-flow* present in each frame of the scene camera video and computed the average *pixel-flow* across all frames of each identified fixation. Then we discarded all fixations containing above average *pixel-flow*. This approach ensures that only true fixations are included in the analysis.

### Bipedal treadmill rig

Locomotion experiments were carried out with a custom-designed primate treadmill rig that enabled animals to perform bipedal locomotion simultaneously with a head-fixed visual task. We constructed the modified primate chair (Rogue Research Inc) with a highly reinforced aluminum frame and stabilized it with weights totaling 300 pounds. This structure was positioned over a modified conveyor belt and equipped with a removable seat. A Faraday cage contained the power supply, voltage regulator, and drive motor to prevent electromagnetic interference with neural recordings. Animals were gradually trained using positive reinforcement. Initially, they could either walk or stand in the rig at will but were encouraged to stand for brief periods with consistent reward of juice or peanuts. Once animals were comfortable standing in the rig, the treadmill motor was turned on at the slowest possible speed (<1 cm/s), and they were given additional rewards to build a positive association with walking. As they became more comfortable with walking in the rig, the speed was gradually increased to the final speed of 8 cm/s. Animals were under constant supervision during recordings and given frequent breaks.

### Locomotion speed estimation

Treadmill velocity was selected based on video recordings of animals walking in a freely-moving arena. Average speed during locomotion was determined by processing video from an overhead camera that captured frames at 30 Hz. First, we used multiple frames to create a composite background image without the animal. As the camera was stationary throughout the recording, the composite background image could be used to extract the animal’s position in each frame. The background frame was subtracted from each frame of the video; then a threshold was used to remove most of the noisy pixels. The pixels that were different from the background image revealed the animal’s shape, and a centroid of this figure was computed to estimate the animal’s position in the environment. The difference in the animal’s position between consecutive frames was used to calculate velocity. To compute an estimate of locomotion velocity, we computed the average speed in two second increments. We next removed periods where the animal was not moving by removing any increments in which the animal’s velocity was less than 10% of the maximum. The average speed when the animal was walking was 5.43 cm/s. In order to maximize changes in arousal levels between the still and walk conditions, a slightly higher treadmill belt velocity of 8 cm/s was used in this study.

### Head-fixed electrophysiology

In the treadmill experiment, electrophysiological data was recorded from multi-electrode arrays using the Cerebus neural signal processor developed by Blackrock Neurotech. The Central software suite (Blackrock Neurotech) recorded putative spike waveforms at 30 kHz based on thresholding of extracellular potentials for each electrode or using the software’s automatic thresholding, which was ±6.25 times the standard deviation of the raw signal. These waveforms were further sorted to isolate single units using Offline sorter (Plexon). Each unit was evaluated as multi- or single-unit using several criteria: consistent spike waveforms, waveform shape (slope, amplitude, trough-to-peak) and exponentially decaying interspike-interval histogram with no interspike interval shorter than the refractory period (1 ms). The analyses here used all single- and multi-unit activity.

### Head-fixed eye tracking

In the treadmill experiment, we measured eye position and pupil diameter using a stationary eye tracker (model ETL-200; ISCAN). This data was recorded at 1 kHz with the neural signal processor (Blackrock Neurotech) which precisely aligned eye and neural data.

### Receptive field mapping of neurons

We identified receptive fields (RFs) of recorded neurons across 24° × 24° of visual space. We first divided this area into a 3×3 grid of nine squares each covering 8° × 8° of visual space, and further divided each of these squares into a 6 × 6 grid with 36 sublocations covering ∼1.33° × ∼1.33° of visual space. In each trial, one of the nine squares was randomly chosen, and a reverse correlation movie of red, blue, green, and white patches was presented in random order across the 36 sublocations. This was repeated for 10 trials in each of the nine squares, making for 90 trials in total. Animals had to maintain fixation during the full duration of each trial. Then, we averaged the responses across all trials to generate receptive field heatmaps for each neuron. Since most recorded units remained stable across days (>70% of units), RF mapping was done prior to the first session and performed once every month during recordings for each monkey.

### Treadmill behavioral task

Animals performed a visual task consisting of 8° visual stimuli flashed over the receptive fields of the recorded neurons on a cathode ray tube (CRT) monitor. In total, there were 8 luminance-matched colored patches, presented 10 times each. Within each trial, stimuli were presented for 448 ms at a viewing distance of 60 cm. Eye movements were monitored throughout the recording session using an infrared eye tracking system (model ETL-200; ISCAN) at a 1-kHz sampling rate. If at any point during the trial, eye position exceeded 1° outside the boundaries of the fixation point (.2°), then the trial was automatically aborted. Otherwise, animals were rewarded with juice at the end of each successful trial. ‘Walk’ and ‘still’ trials occurred in a blocked manner in random order for each session; we did not observe significant changes in neural responses in any area depending on which block occurred first (P > 0.1, Wilcoxon signed-rank test). Stimulus presentation was recorded and synchronized with the neural data using a custom-designed experimental control unit.

#### Visual stimuli

Selected colors were isoluminant and equally distributed across the CIELUV color space, which is perceptually uniform. These color stimuli were presented on an isoluminant grey background with Psychophysics Toolbox. Chromaticity and luminance were calibrated for the colors and grey background using a Tektronix photometer (J17 LumaColor, Tektronix). Find RGB and LUV values for the color stimuli below:

| R | G | B | L | U | V |
|---|---|---|---|---|---|
| 147 | 75 | 4 | 39.99 | 58.50 | 36.68 |
| 81 | 99 | 0 | 39.02 | -8.17 | 44.27 |
| 0 | 100 | 83 | 37.37 | -30.58 | 7.18 |
| 0 | 96 | 91 | 36.22 | -28.51 | -2.02 |
| 0 | 95 | 101 | 36.22 | -27.40 | -10.96 |
| 138 | 20 | 201 | 38.06 | 25.45 | -99.69 |
| 214 | 55 | 155 | 51.15 | 87.74 | -38.09 |
| 226 | 58 | 129 | 52.71 | 106.54 | -13.64 |

#### Freely moving color change task

The experimental setup consisted of a custom-made cage (120 cm × 60 cm × 90 cm) located in a dedicated room, free from distractions. Monkeys were habituated to the environment at least 4 days per week for over 4 weeks. During this time, they were trained to press the button on each box to receive a reward. After completing training, monkeys were tested with a range of balanced and unbalanced reward schedules. For balanced schedules, both stations were set to a reward schedule of 20 s or 30 s. For unbalanced schedules, we used combinations such as 20 s on one station and 40 s on the other; other unbalanced schedules included 15 s vs. 25 s and 10 s vs. 30 s. During some sessions, the unbalanced schedules were reversed one to three times: for example, a station initially set to 20 seconds would switch to 40 seconds and vice versa. Each session lasted until the monkey earned 100 or 200 rewards, taking between 1 - 2 hours, including a 1-hour break after 100 rewards in the longer sessions. If monkeys were not engaged with the task for more than 2 min, we occasionally intervened to encourage them to re-engage with the task.

The stations initially displayed a blue screen, indicating that the reward was not available. Then, during each time bin of size dt = 10 ms, a sample was drawn from a Bernoulli distribution with probability of dt/VI. If the sample drawn was 0, then the screen remained blue and another sample was drawn in the next time bin. Conversely if the sample drawn was 1, then the screen would turn red, indicating that reward was now available, and remain red until the animal collected the reward. Overall, this resulted in an exponential probability function that asymptotically approached 1 over time, but at different rates depending on the VI schedule.

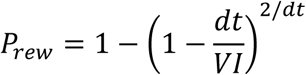

### Electrical stimulation

Stimulation experiments were conducted using a RZ5/IZ2-32H electrical stimulation system (Tucker Davis Technologies). The RZ5 stimulator was connected to a head-stage Y-adaptor (Blackrock Neurotech), which allowed for simultaneous recording and stimulation. Each V4 or dlPFC channel received 10 trials of stimulation, consisting of 5 biphasic current pulses (0.2 ms pulse duration, 100 μA, 50 Hz), with a 5 s delay period between trials. We stimulated all available 32 channels in both V4 and dlPFC, one at a time. Then, we examined the firing rates across all neurons in the target population (V4 or dlPFC) in a given session and identified peaks in the activity corresponding to the stimulation period. Finally, we used Wilcoxon rank-sum tests to compare evoked neural responses in the 100-ms following the generation of the first pulse between the Still and Walk conditions relative to the ongoing activity in the 100-ms interval before the first electrical pulse.

### Subspace analysis

We performed a communication subspace analysis as previously described^10^, assessing communication both within and between brain regions. Briefly, we computed the residuals of each neuron by subtracting its mean response to obtain trial-by-trial fluctuations around the mean (**Fig. 3a**). Next, we partitioned our dataset into a training dataset and testing dataset. We fit a linear regression to the training set, using the fluctuations in one neural population to predict the fluctuations in another. This regression takes the form *Y* = *XB*, where *B* assigns a “weight” to each neuron in population 1 to predict the activity of each neuron in population 2. Next, we applied this model to the test dataset, and used the activity in population 1 to make predictions about what the neural activity we should expect in population 2 (**Fig. 3b**). Finally, we compared these predictions to the actual recorded activity in population 2. The accuracy of this model provided a measure of communication between the two neural populations. Importantly, this analysis can be used to selectively examine feedforward or feedback interactions between areas by shifting spike trains by the relevant time delay. Here, we shifted dlPFC neural activity by ±50 ms relative to V4 activity to examine feedforward or feedback interactions respectively (**Fig. 3c**). Furthermore, in order to assess the dimensionality of this communication, we used a reduced rank linear regression, which imposes a rank constraint on the matrix *B*. By using only the top principal components of *B*, we can see how many dimensions are necessary to account for communication within and between brain regions. The dimensionality of this communication can be defined as the number of dimensions needed for the prediction accuracy to plateau.

### Spike train cross-correlation analysis

To measure the synchrony in spiking activity between neuron pairs – both within and across brain areas – at the time scale of milliseconds, we computed their spike train cross-correlograms^19,20^. We represented the spike train of each neuron during the fixation period as a binary time series with 1ms resolution – each data point of the series was either 1 if the neuron fired an action potential during the corresponding 1ms window or 0 otherwise. Trial averaged cross correlations of the spike trains between neurons j and k were computed as:

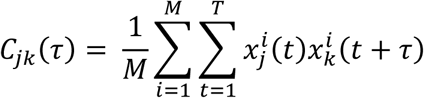

where M is the number of trials, T is the duration of the spike train segments, x is the neural response and τ is the lag. We also corrected the CCG to remove any stimulus induced correlations by subtracting a shuffle predictor – efficiently computed from the cross-correlation of the post-stimulus time histograms (PSTHs)^21–24^. For investigating coupling of V4 and dlPFC neurons, we looked at CCGs that had a significant peak at lags between 30 to 50 ms (positive and negative) that were larger than 2 times the standard deviation. By taking the area under the curve of the cross-correlogram at the relevant time delay (±15 ms around the peak), we computed the synaptic efficacy between neuron pairs.

### Convolutional Autoencoder (CA)

We trained a convolutional autoencoder (CA) to compress images from the scene camera into concise, 128-dimensional codes. We made sure that we were not overfitting to the data by monitoring the reconstruction error on a held-out set of images, and stopped training at 300 epochs when the reconstruction error plateaued. We then applied PCA to the 128-D vector representations to further compress each image to a 16-dimensional vector representation. The Euclidean distance between two vectors was used to quantify their visual similarity. We used a k-d tree data structure to organize the vectors and quickly search for the nearest neighbors of any given frame ^25^.

Clusters of similar fixations were created by randomly choosing a seed vector and finding its ‘k’ nearest (Euclidean distance) neighbors in PCA space. All fixations belonging to a cluster had similar visual content. When creating a pair of clusters, if some fixations belonged to both clusters, they were randomly assigned to only one of them.

### d-prime

To assess the discriminability of neural responses between different stimuli, we computed d-prime (d′) following established methods in neuroscience. *d-*prime is a widely used measure that quantifies the signal-to-noise ratio in a discrimination task, indicating the ability of a neuron to distinguish between two classes of stimuli. For each neuron in each session, the mean responses (*u*_A_ and *u*_B_) to stimuli A and B as well as the standard deviations of responses were obtained. We then computed *d*′ using the following formula:

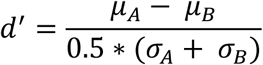

Here, *u*_A_ and *u*_B_ are mean responses of one neuron to stimuli A and B, respectively. Then computed this value for all adjacent stimuli across the tuning curve of 8 color patches.

### Statistics

For the pupil analysis, we computed average pupil size, velocity and position variance. Then a Wilcoxon rank-sum test was used to compare these values across the still and walk conditions, as well as across different locomotion speeds.

Except otherwise indicated, firing rates were smoothed with a Gaussian kernel with a standard deviation of 50 ms. For the population analysis in **Fig. 1e**, we took each neuron’s average firing rate during the stimulus period and used a Wilcoxon rank-sum test to compare between the still and walk conditions. Here, we corrected for multiple comparisons using the false discovery rate, which has a standardized implementation in MATLAB Finally, for the grand average of all neurons in each brain area, we used a Wilcoxon rank-sum test to compare the average firing rate of the entire neural population between the still and walk conditions.

In the spike timing coordination analysis, we used the raw rasters to compute the cross-correlograms between neuron pairs both within and across brain regions. We added a shuffle correction to account for changes in firing rate. Then we computed the area under the curve of the cross-correlogram at the relevant time delay (±15 ms around the peak), excluding any timepoints that were below chance. Finally, a paired t-test was used to compare areas under the curve between the still and walk conditions, subsampled 10 times at n = 500 neuron pairs.

For the subspace analysis, residuals were computed by subtracting the average stimulus response from each trial to obtain the trial-by-trial fluctuations. In the locomotion paradigm, average stimulus responses were computed separately for each of the 8 stimuli. Meanwhile, for the freely-moving data, average stimulus responses were computed across all the fixations in the dataset. This dataset was divided into a training and testing dataset using a cross-validation procedure with 10 folds for each session. For within area predictions, half the neural population was used to predict the other half, after shuffling the neurons to prevent oversampling on adjacent channels. For between-area predictions, the fluctuations in one area were used to predict fluctuations in another at a 50 ms delay. Then, we compared the predicted fluctuations to the actual ones, and computed a mean squared error. We divided this error by the chance mean squared error to obtain the normalized mean squared error: chance error was defined as the mean squared error when we predict the average value for every timepoint in the test data. Finally, we defined accuracy as 1 minus the normalized mean squared error, such that 1 represented perfect accuracy and 0 represented chance accuracy. Throughout this analysis, we used a reduced-rank linear regression, varying the rank constraint from 1 to 10. The dimensionality of communication could be defined as the number of dimensions needed for the prediction accuracy to plateau. Negative prediction accuracies were zeroed out, and then a t-test was used to compare prediction accuracies between the still and walk conditions.

Finally, in the electrical stimulation paradigm, firing rates were smoothed with a moving mean of 25 ms to retain finer temporal resolution. Then, we computed the average responses across the entire dlPFC population for each stimulated electrode in V4. In the population average, we found peaks in the still and walk conditions during the electrical stimulation period. Next, we used a two-sample t-test to compare the height of these peaks across the still and walk conditions. Again, we corrected for multiple comparisons using the false discovery rate.

---

**Extended Data Figure 1.**
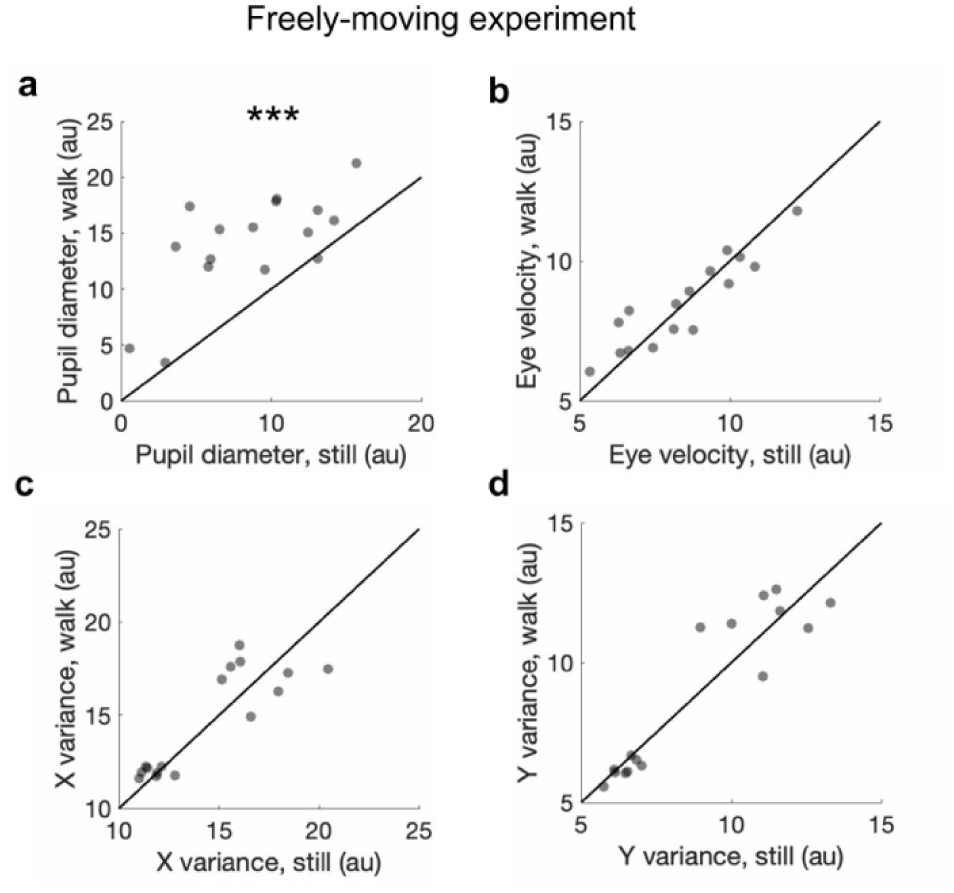
**a,** Pupil diameter during fixations across the still and walk conditions (Wilcoxon signed-rank test; p = 2.23 × 10^-5^, n = 16 sessions). b, Eye velocity during fixations across still and walk conditions (Wilcoxon signed-rank test, p = 0.79, n = 16 sessions). c, Variance of eye X-position across the still and walk conditions (two-sample t-test, p = 0.66, n = 16 sessions). d, Variance of eye Y-position across the still and walk conditions (two-sample t-test, p = 0.94, n = 16 sessions). Each dot represents a session. * p < 0.05. ** p < 0.01, *** p < 0.001.

**Extended Data Figure 2.**
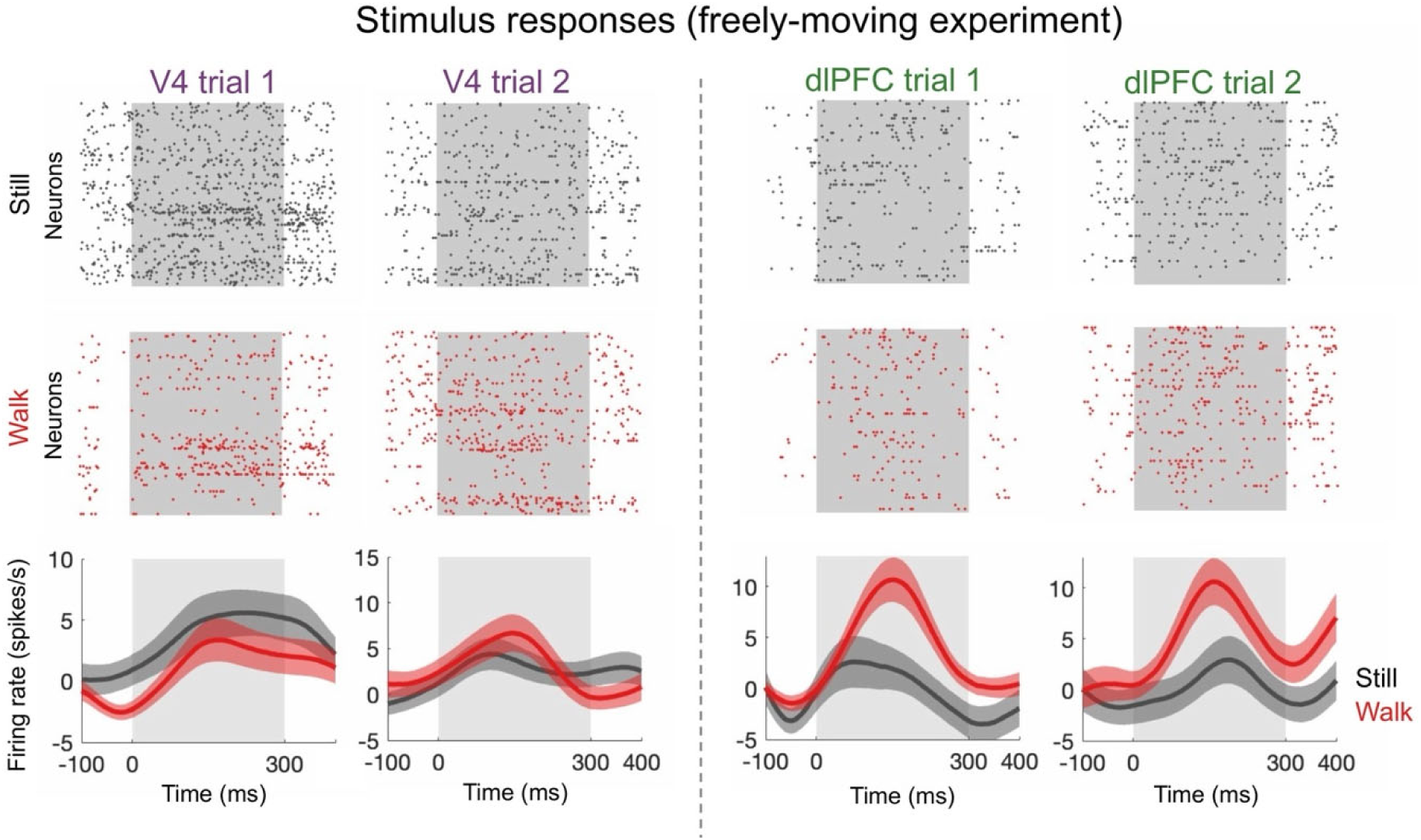
Example stimulus responses to visual fixations in V4 and dlPFC during the still and walk conditions in the freely-moving experiment. Traces show mean +/- SEM.

**Extended Data Figure 3.**
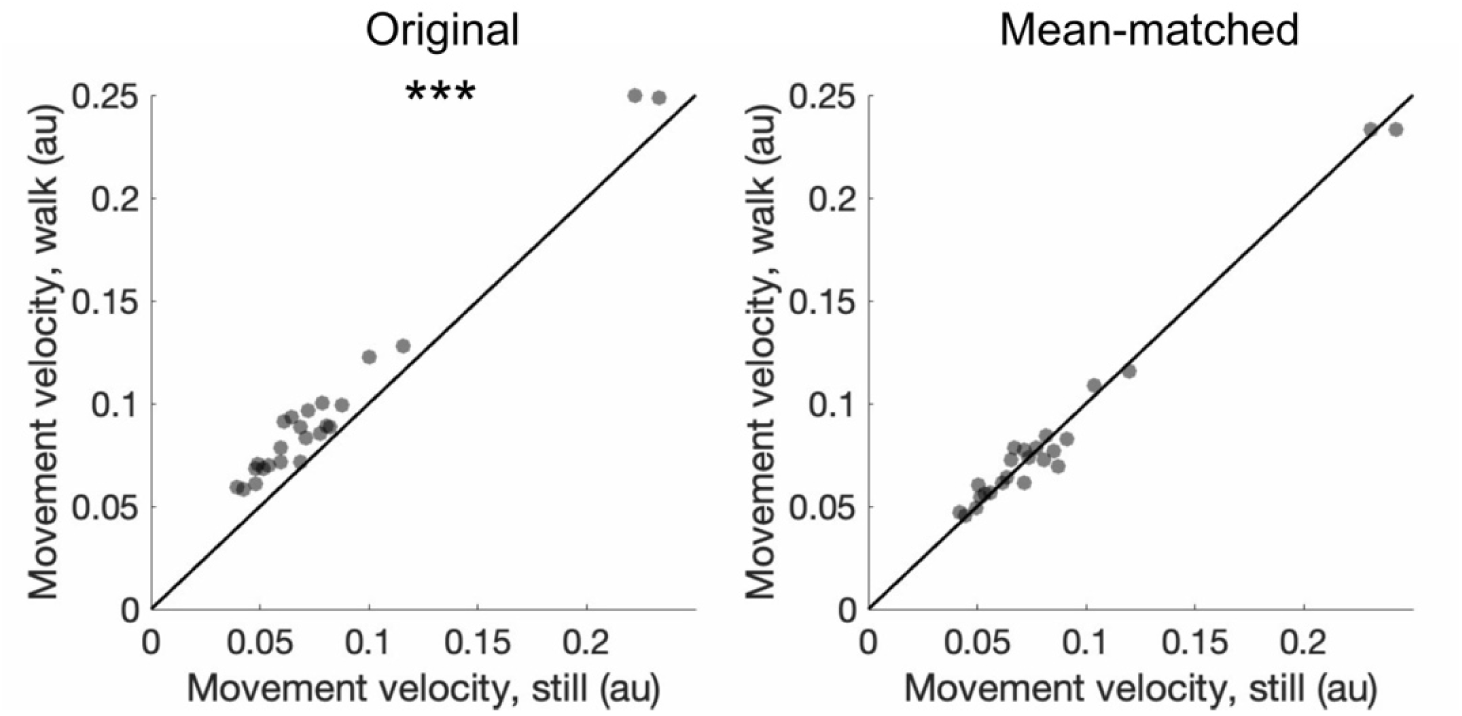
Left panel: movement velocities in the original data during the still and walk conditions (one-way t-test, p = 3.10 × 10^-11^, n = 25 sessions). Right panel: movement velocities after mean-matching during the still and walk conditions (one-way t-test, p = 0.86, n = 25 sessions); 10% of fixations were excluded. Each dot represents a session. * p < 0.05. ** p < 0.01, *** p < 0.001.

**Extended Data Figure 4.**
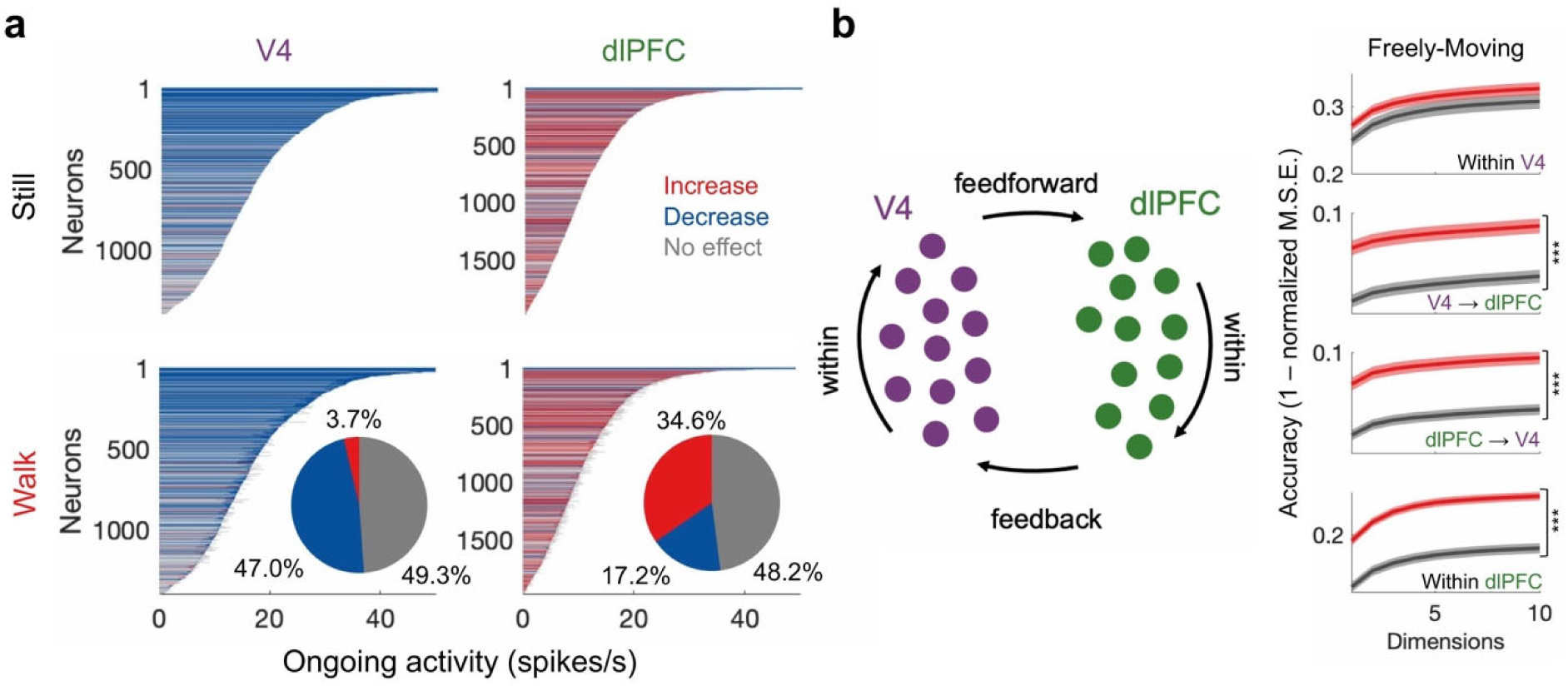
**a,** Stimulus responses in V4 (Wilcoxon rank-sum; p = 1.43 × 10^-151^, n = 1,395 neurons) and dlPFC (Wilcoxon rank-sum test; p = 4.12 10^-27^, n = 1,979 neurons) sorted based on response magnitude for movement-matched fixations across the still and walk conditions. Inset shows proportion of neurons that increased, decreased, or maintained their firing rates during locomotion. **b,** Subspace communication for movement-matched fixations across the still and walk conditions. Traces show mean +/- SEM. * p < 0.05. ** p < 0.01, *** p < 0.001.

**Extended Data Figure 5.**
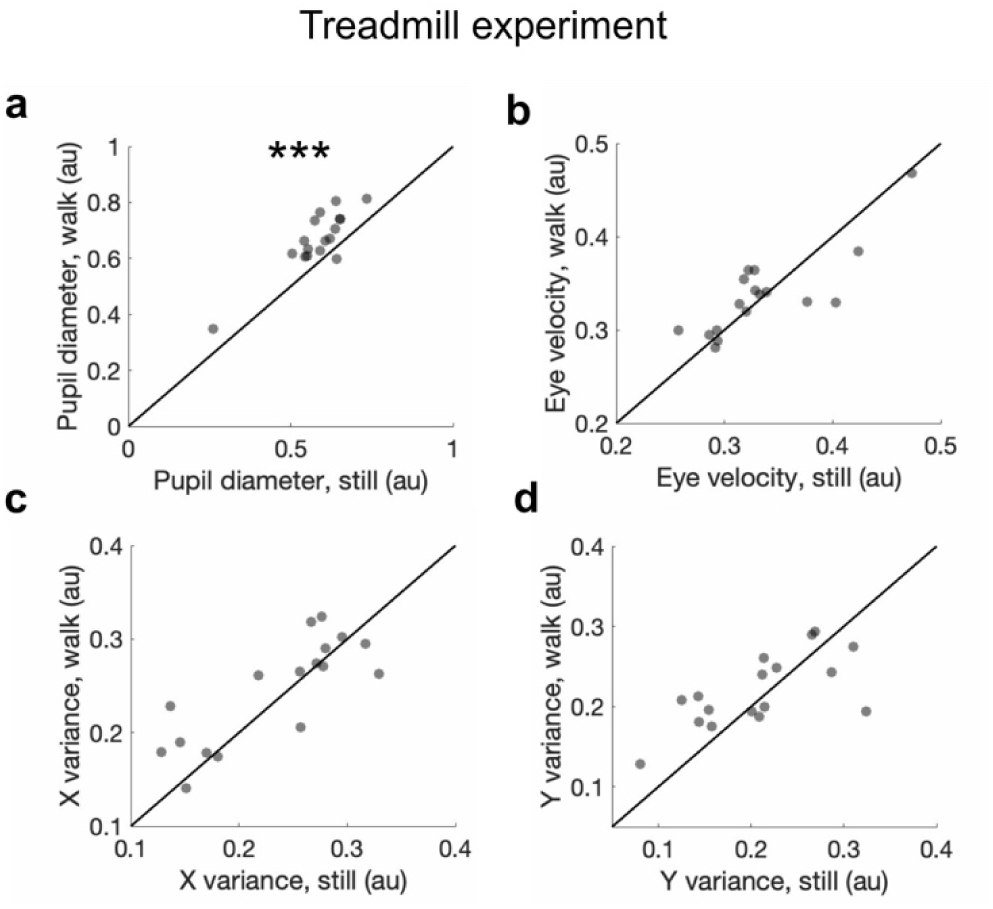
**a,** Pupil diameter during fixations in the still and walk conditions (Wilcoxon signed-rank test; p = 5.58 × 10^-7^, n = 17 sessions). **b,** Eye velocity during fixations in the still and walk conditions (Wilcoxon signed-rank test, p = 0.83, n = 17 sessions). **c,** Variance of eye X-position during fixations in the still and walk conditions (two-sample t-test, p = 0.24, n = 17 sessions). **d,** Variance in eye Y-position during fixations in the still and walk conditions (two-sample t-test, p = 0.37, n = 17 sessions). Each dot represents a session. * p < 0.05. ** p < 0.01, *** p < 0.001.

**Extended Data Figure 6.**
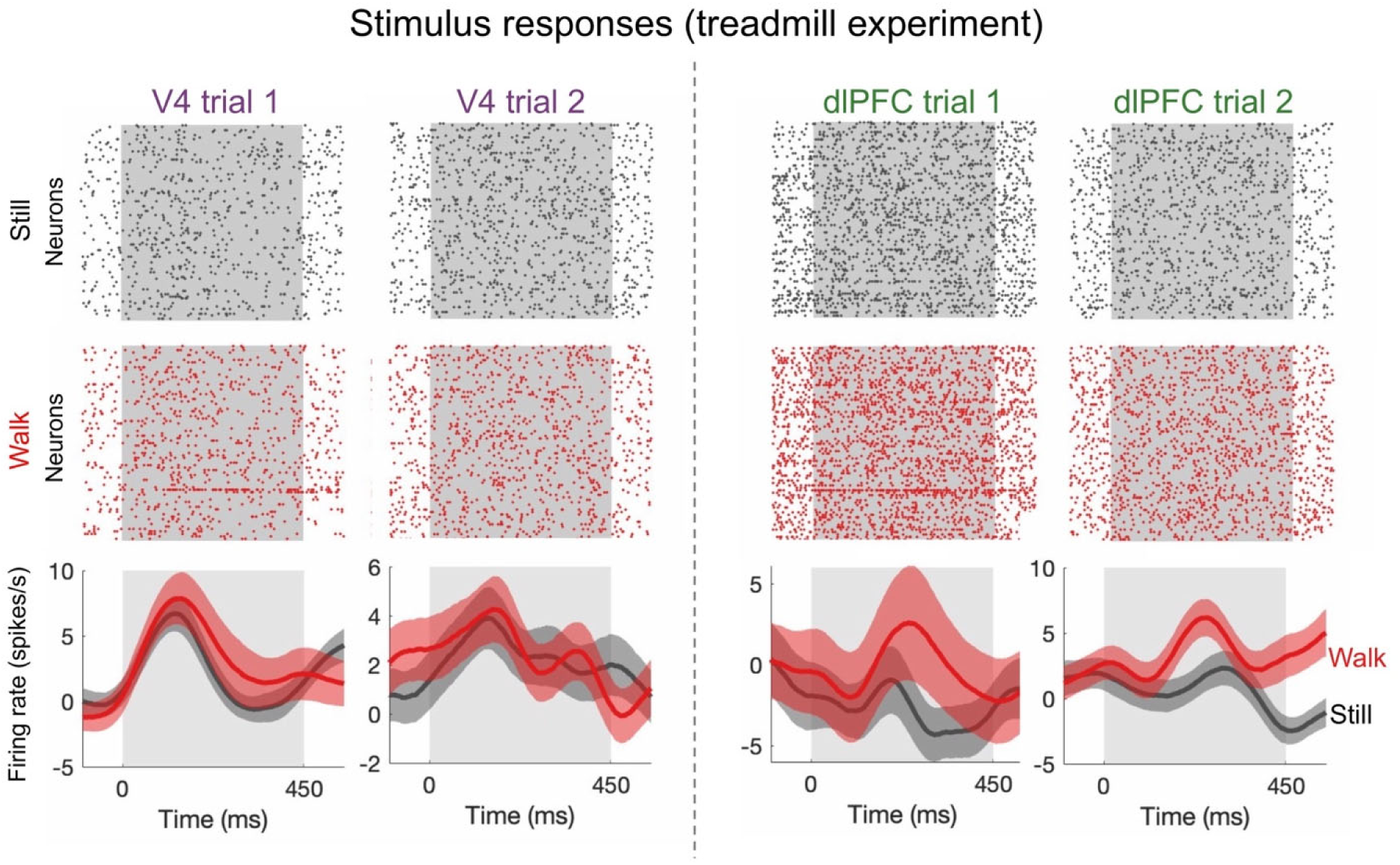
Example stimulus evoked responses in V4 and dlPFC during the still and walk conditions in the treadmill experiment. Traces show mean +/- SEM.

**Extended Data Figure 7.**
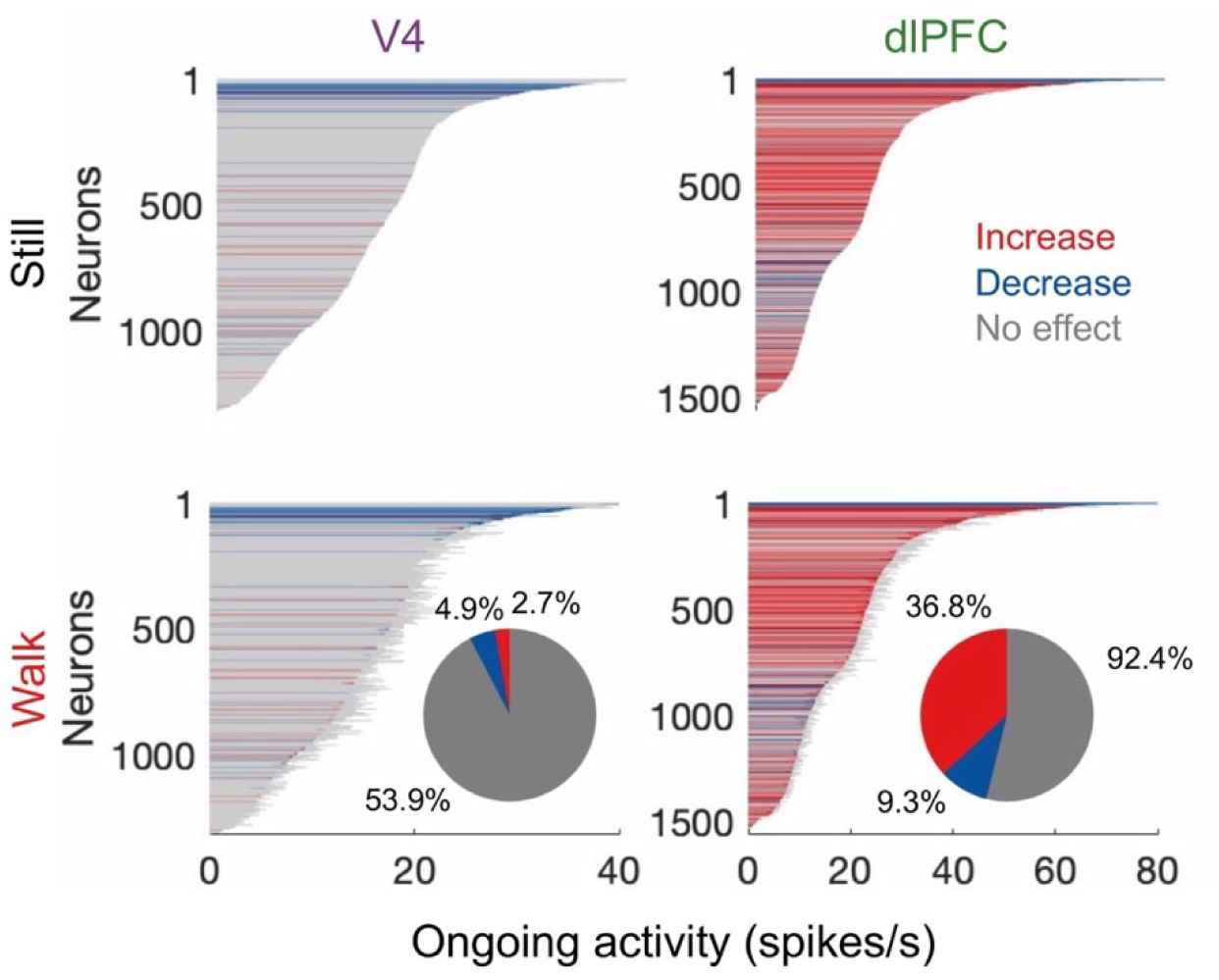
Ongoing activity for neurons recorded in the treadmill experiment in V4 (no significant effect, Wilcoxon signed-rank test: p = 0.59, n = 1,315 neurons) and dlPFC (12.61% increase, Wilcoxon signed-rank test: p = 1.01 × 10^-66^, n = 1,558 neurons). Inset shows proportion of neurons that increase, decrease, or maintain their firing rates during locomotion. Red: neurons that significantly increase their firing rates during walk, blue: neurons that decrease their firing rates, gray: neurons with no significant change.

**Extended Data Figure 8.**
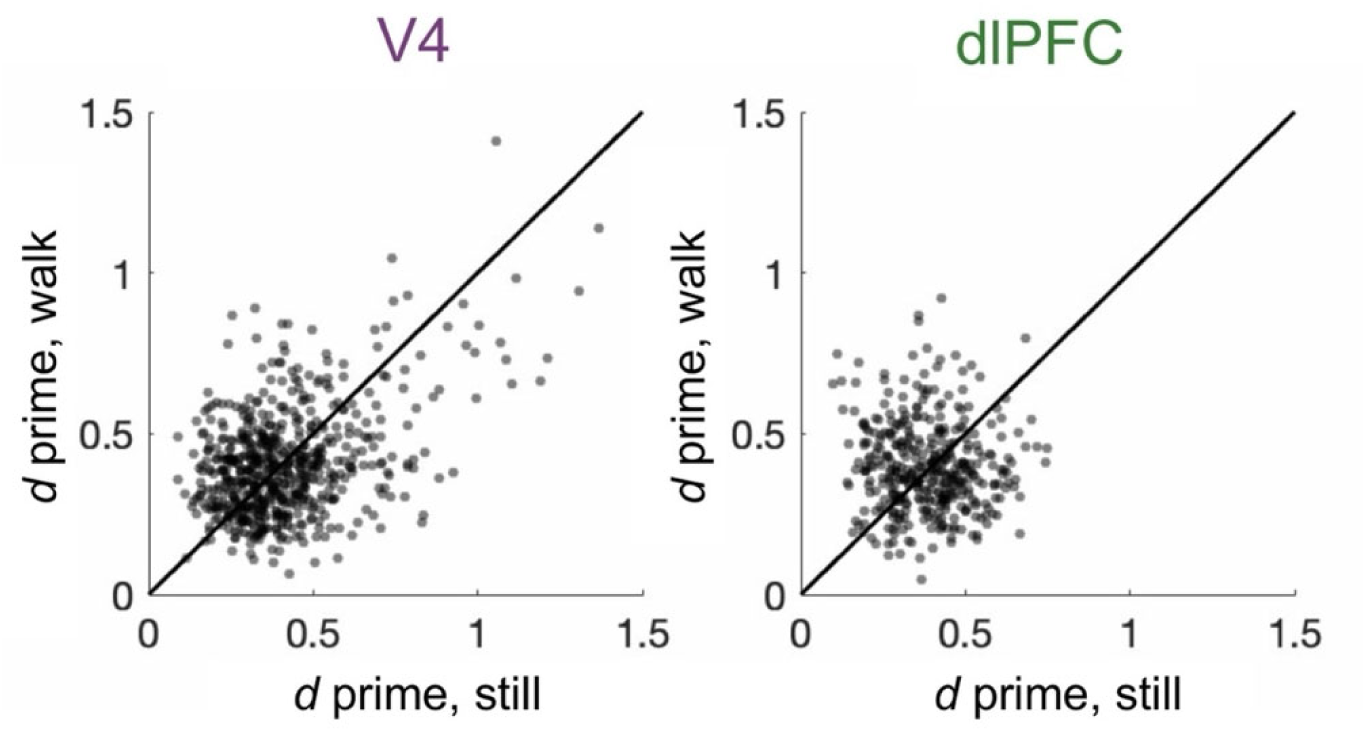
Selectivity of stimulus-responsive neurons in V4 (two-sample t-test: p = 0.50, n = 628 neurons) and dlPFC (two-sample t-test: p = 0.40, n = 389) across the still and walk conditions of the treadmill experiment. Here, each dot represents a neuron. * p < 0.05. ** p < 0.01, *** p < 0.001.

**Extended Data Figure 9.**
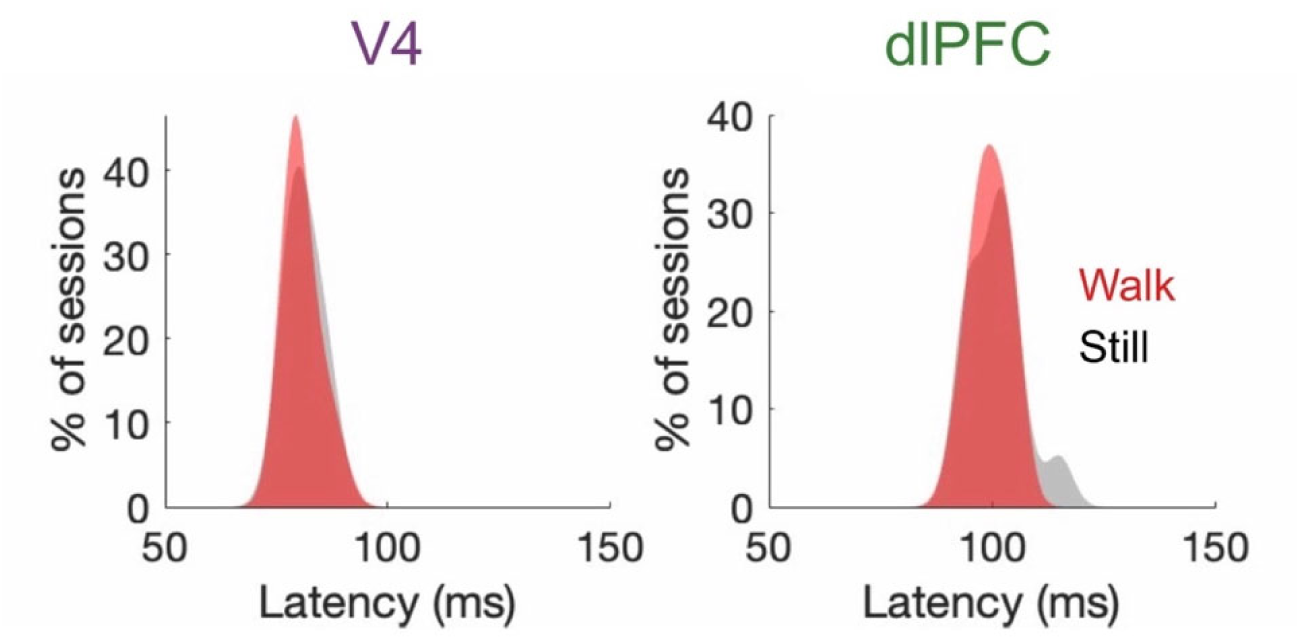
Response latency (time to reach half peak firing rate) for V4 (two-sample t-test: p = 0.79, n = 13 sessions) and dlPFC neurons (two-sample t-test: p = 0.52, n = 13 sessions) across the still and walk conditions of the treadmill experiment. Mean latency is presented across sessions. * p < 0.05. ** p < 0.01, *** p < 0.001.

**Extended Data Figure 10.**
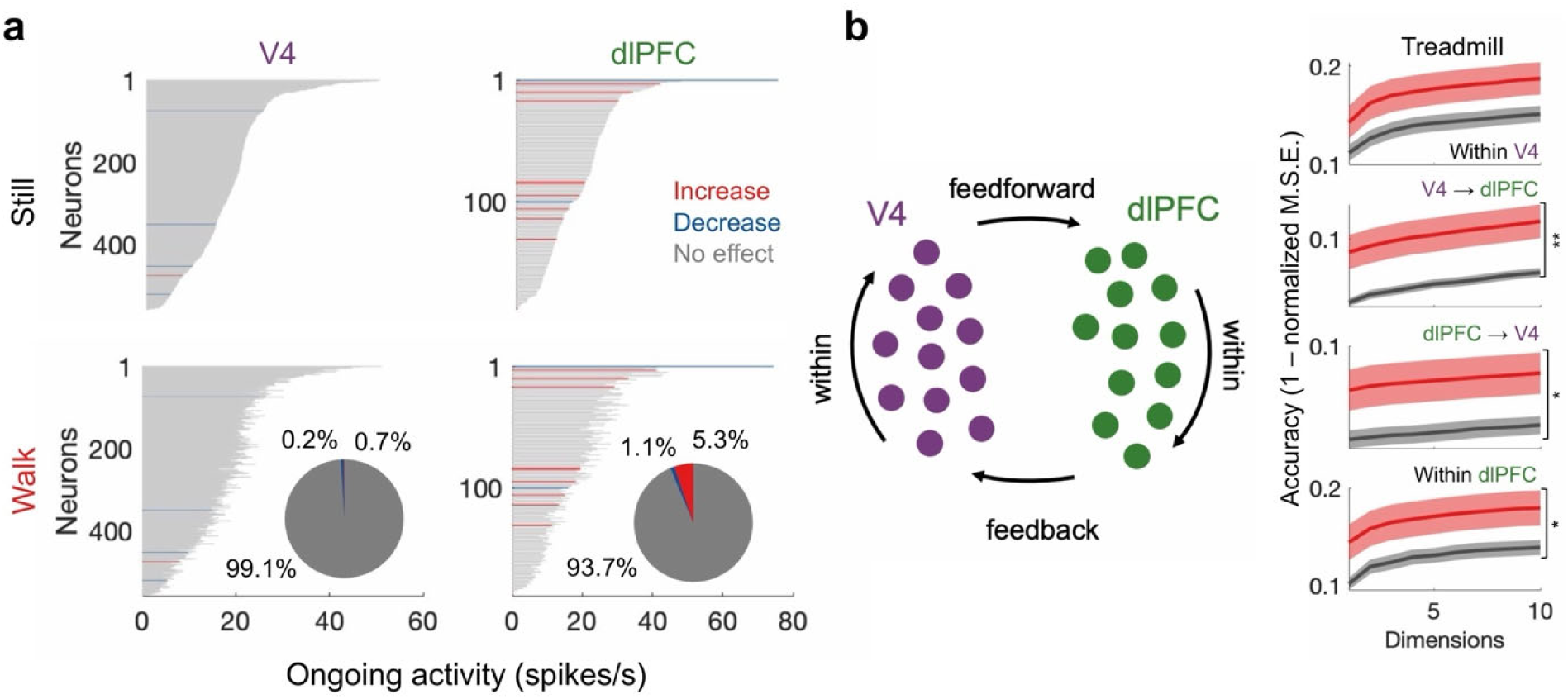
**a,** Stimulus responses in V4 (Wilcoxon signed-rank test: p = 0.033, n = 559 neurons) and dlPFC (Wilcoxon signed-rank test: p = 5.00 10^-6^, n = 190 neurons) sorted based on response magnitude after mean-matching neuronal responses (no change in ongoing firing rate between still and walk conditions). Inset shows proportion of neurons that increased, decreased, or maintained their firing rates during locomotion. **b,** Subspace communication for mean-matched neurons that did not change their ongoing firing rates between the still and walk conditions. Traces show mean +/- SEM. * p < 0.05. ** p < 0.01, *** p < 0.001.

**Extended Data Figure 11.**
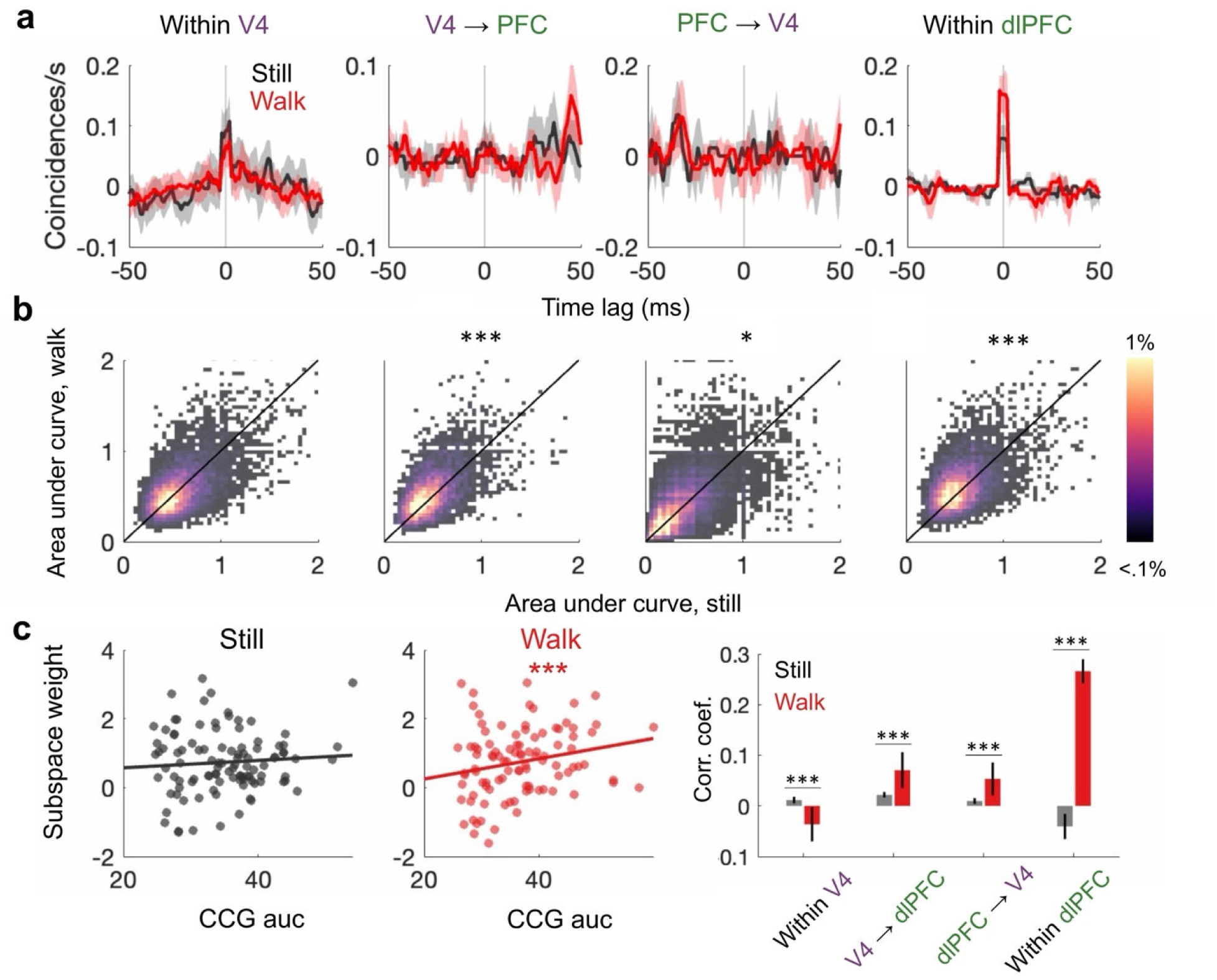
**a,** Example cross-correlograms both within and between brain regions for the still and walk conditions. Traces show mean +/- SEM. **b,** Population spike-time coordination both within and between brain areas during the still and walk conditions. This analysis revealed that locomotion caused a significant increase in spike-time coordination between areas (V4 to dlPFC: Wilcoxon signed-rank test; p = 9.04 × 10^-4^, 38,955 neuron pairs subsampled at n = 500; dlPFC to V4: Wilcoxon signed-rank test; p = 0.0241, 38,955 neuron pairs subsampled at n = 500) as well as within dlPFC (Wilcoxon signed-rank test; p = 4.56 × 10^-4^, n = 30,588 neuron pairs subsampled at n = 500). However, locomotion did not modulate spike-time coordination within V4 (Wilcoxon signed-rank test; p = 0.37, 29,948 neuron pairs subsampled at n = 500). This suggests that brain-state modulates not only firing rates but also interareal communication in a region-specific manner. **c,** Left panel: correlation between area under the curve in the cross-correlogram analysis and the regression weights in the subspace analysis. Right panel: correlation between CCG and subspace weights both within and between brain regions. Here, we found a significant positive correlation between subspace weights and CCG within V4 (one-sample t-test, still: p = 0.014, 1,316 neurons subsampled 100 times at n = 250), from V4 to dlPFC, (one-sample t-test, still: p = 0.0011, walk: p = 0.0013, 1,316 neurons subsampled 100 times at n = 250), from dlPFC to V4 (one-sample t-test, still: p = 0.061, walk: p = 1.11 × 10^-16^, 1,865 neurons subsampled 100 times at n = 250), and within dlPFC (one-sample t-test, walk: p = 0.20, 1,316 neurons subsampled 100 times at n = 250). Meanwhile, two regions showed negative correlations between subspace weights and CCG: within V4 (one-sample t-test, walk: p = 0.0028, 1,316 neurons subsampled 100 times at n = 250) and within dlPFC (one-sample t-test, still: p = 0.0015, 1,316 neurons subsampled 100 times at n = 250). Importantly, we found that walking modulated the correlation between subspace weights and CCG, increasing the correlation from V4 to dlPFC (two-sample t-test: V4 to dlPFC: p = 0.024, n = 1,316 neurons), from dlPFC to V4 (two-sample t-test: V4 to dlPFC: p = 0.0045, n = 1,316 neurons) and within dlPFC (two-sample t-test: p = 3.86 × 10^-15^, n = 1,865 neurons), but decreasing it within V4 (two-sample t-test: 1.33 × 10^-4^, n = 1,316 neurons). This suggests that brain-state can systematically shifts communication to neural pairs with higher or lower synaptic strength. Error bars show SEM. * p < 0.05. ** p < 0.01, *** p < 0.001.

**Extended Data Figure 12:**
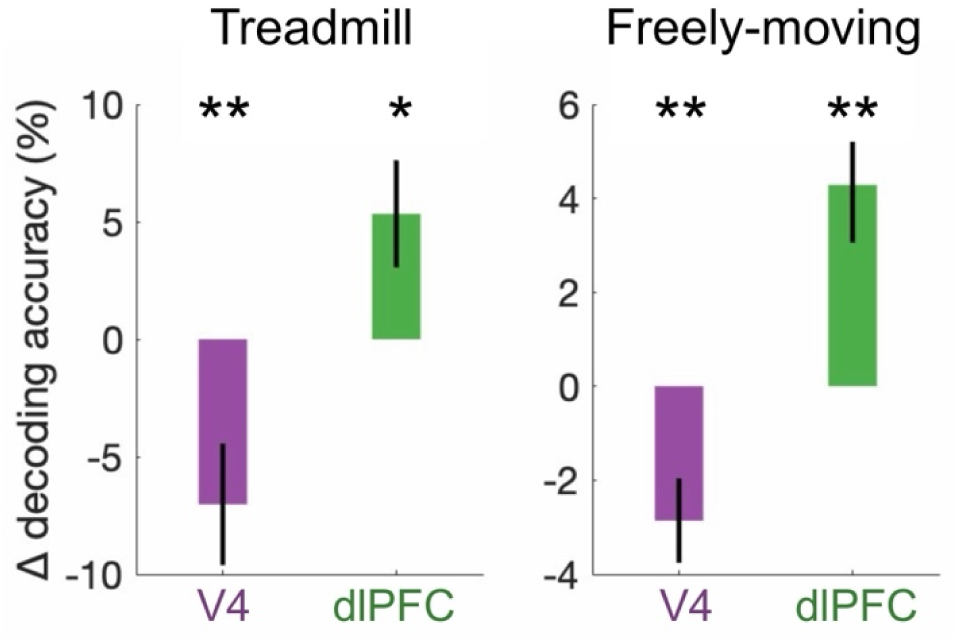
Change in stimulus decoding accuracy based on population activity across the still and walk conditions (relative to chance level). In both the treadmill and freely-moving experiments, locomotion significantly decreased decoder accuracy in V4 (treadmill: p = 0.010, n = 130 folds; freely-moving: p = 0.0082, n = 27 sessions) but increased it in dlPFC (treadmill: p = 0.03, n = 130 folds; freely-moving: p = 0.0027, n = 27 sessions). Here, we tested the relative change in decoding for each session, rather than the absolute value.

**Extended Data Figure 13.**
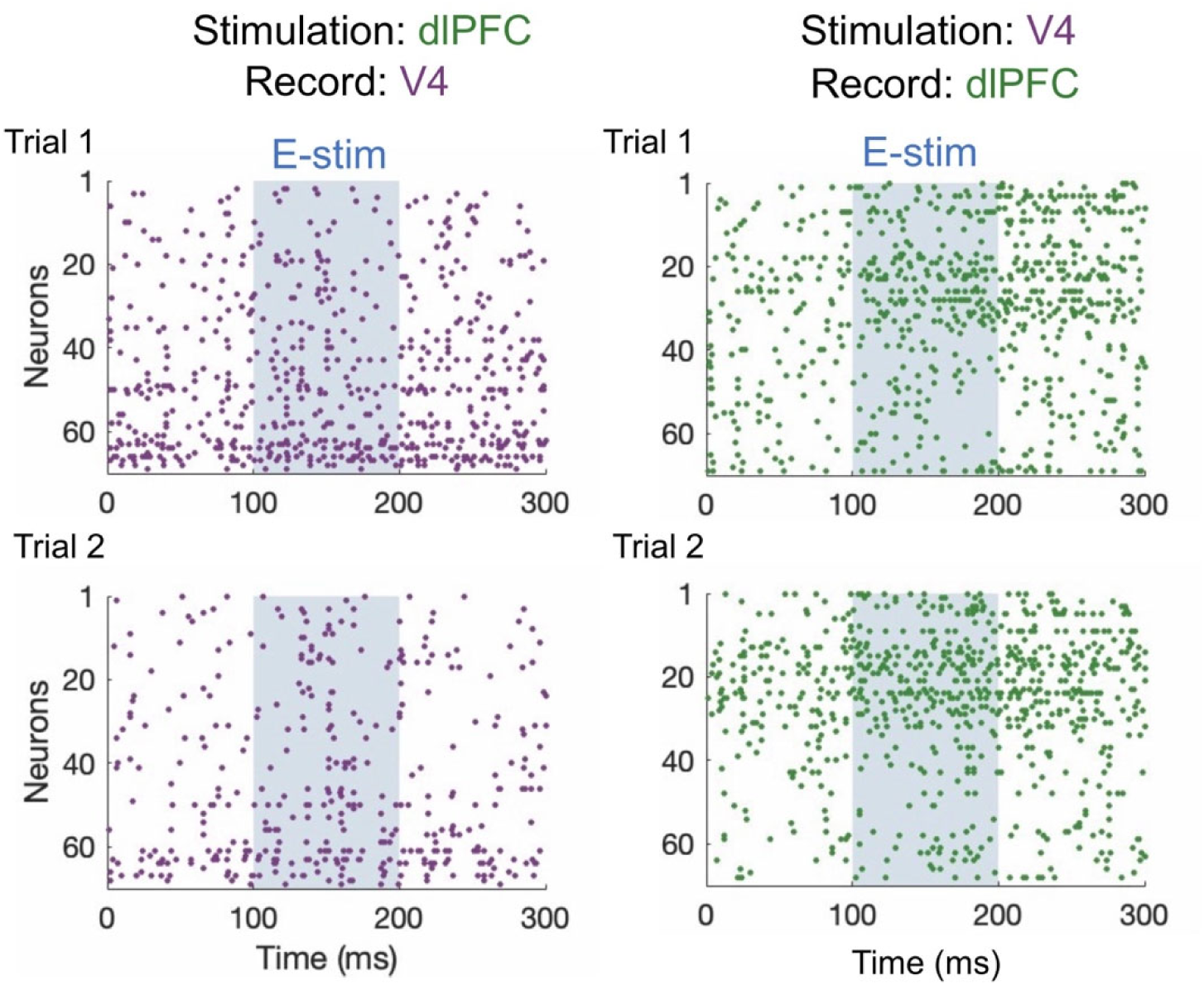
Example evoked responses in dlPFC/V4 to electrical stimulation in V4/dlPFC during the walk condition. Rasters represent the spiking activity of each neuron on the array in one trial. Left: electrical stimulation in dlPFC and recording in V4; right: electrical stimulation in V4 and recording in dlPFC. Blue shaded regions represent the electrical stimulation interval.

